# Loss of *RREB1* reduces adipogenesis and improves insulin sensitivity in mouse and human adipocytes

**DOI:** 10.1101/2024.07.30.605923

**Authors:** Grace Z. Yu, Nicole A. J. Krentz, Liz Bentley, Meng Zhao, Keanu Paphiti, Han Sun, Julius Honecker, Marcus Nygård, Hesam Dashti, Ying Bai, Madeleine Reid, Swaraj Thaman, Martin Wabitsch, Varsha Rajesh, Jing Yang, Katia K Mattis, Fernando Abaitua, Ramon Casero, Hans Hauner, Joshua W Knowles, Joy Y Wu, Susanne Mandrup, Melina Claussnitzer, Katrin J Svensson, Roger D. Cox, Anna L. Gloyn

## Abstract

There are multiple independent genetic signals at the *Ras-responsive element binding protein 1* (*RREB1*) locus associated with type 2 diabetes risk, fasting glucose, ectopic fat, height, and bone mineral density. We have previously shown that loss of *RREB1* in pancreatic beta cells reduces insulin content and impairs islet cell development and function. However, RREB1 is a widely expressed transcription factor and the metabolic impact of RREB1 loss *in vivo* remains unknown. Here, we show that male and female global heterozygous knockout (*Rreb1*^+/-^) mice have reduced body length, weight, and fat mass on high-fat diet. *Rreb1^+/-^* mice have sex- and diet-specific decreases in adipose tissue and adipocyte size; male mice on high-fat diet had larger gonadal adipocytes, while males on standard chow and females on high-fat diet had smaller, more insulin sensitive subcutaneous adipocytes. Mouse and human precursor cells lacking RREB1 have decreased adipogenic gene expression and activated transcription of genes associated with osteoblast differentiation, which was associated with *Rreb1*^+/-^ mice having increased bone mineral density *in vivo*. Finally, human carriers of *RREB1* T2D protective alleles have smaller adipocytes, consistent with RREB1 loss-of-function reducing diabetes risk.

## Main

Diabetes is a metabolic disease characterized by hyperglycemia due to defects in insulin secretion and/or function. Multiple genetic association signals have been identified in the *Ras-responsive element binding protein 1* (*RREB1*) locus that alter risk for T2D^1–3^ and other metabolic and anthropomorphic traits, including fasting glucose^1,4^, visceral adiposity^5^, waist-hip ratio^6^, bone mineral density^7^, and height^8^. Fine-mapping studies demonstrated that a nonsynonymous variant (p.Asp1171Asn)^9^ is responsible for one of the T2D association signals, supporting *RREB1* as the effector transcript at the locus. RREB1 is a zinc finger transcription factor that binds to RAS-responsive elements found in gene promoters, acting as both a transcriptional activator^10,11^ and repressor^12,13^. Microdeletions on chromosome 6p, which includes *RREB1* in the minimal region, have been identified as a cause of Noonan Syndrome, characterized by short stature and heart abnormalities^14^. Previous studies have also identified a role for Rreb1 in epithelial-to-mesenchymal transition during early embryogenesis^15^, brown fat adipogenesis^16^ and in muscle cells^17^.

As *RREB1* is widely expressed^18^ and there are genetic associations at this locus with multiple phenotypes and traits, it is unclear in which tissues *RREB1* perturbation contributes to diabetes risk. Multi-omic integration of genetic, transcriptomic, and epigenomic data supports equal contribution of islet, adipose, liver, and muscle in mediating disease risk^19^. We have previously found that *RREB1* knockdown or knockout negatively impacts beta cell development and function^20^, suggesting that T2D-risk alleles result in a loss-of-function. Additionally, our previous findings in zebrafish suggest that loss of Rreb1 has effects in other metabolic tissues^21^. To understand the relevant contributions of different tissues on the genetic association of *RREB1* with T2D and related metabolic traits including adiposity, we characterized a global Rreb1 heterozygous knockout mouse model and investigated the effects of *RREB1* perturbation on human adipocyte development and function.

## Results

### Global heterozygous *Rreb1* knockout mice have reduced length, body weight, and fat mass

To explore the impact of RREB1 loss on metabolic traits, we generated global heterozygous knockout (*Rreb1*^+/-^) mice as homozygous null mice are embryonic lethal^14,22^. Global *Rreb1^+/-^* mice were placed on one of three diets at the time of weaning: a high-fat diet (HFD: 60% fat, 20% protein, 20% carbohydrates); calorie-matched low-fat diet (LFD: 10% fat, 20% protein, 70% carbohydrates); or SDS rat and mouse No.3 Breeding diet (RM3: 11% fat, 27% protein, 62% carbohydrates). Male *Rreb1^+/-^* mice had reduced body length on HFD at 29 weeks of age (Fig. 1a). However, there were no significant differences in length for male *Rreb1^+/-^* mice on LFD at 29 weeks of age (Fig. 1a) or at 38 weeks of age on an RM3 diet (Fig. 1b), suggesting that the effect on body length is dependent on calorie intake. Female *Rreb1^+/-^* mice were significantly smaller in length than control mice on HFD at 29 weeks of age (Fig. 1c), but showed a trend on RM3 diet (*p*=0.068) at 38 weeks of age (Fig. 1d). These results are consistent with the genetic association of variation at the *RREB1* locus with height and our previous observations in zebrafish^21,23,24^.

**Fig. 1.**
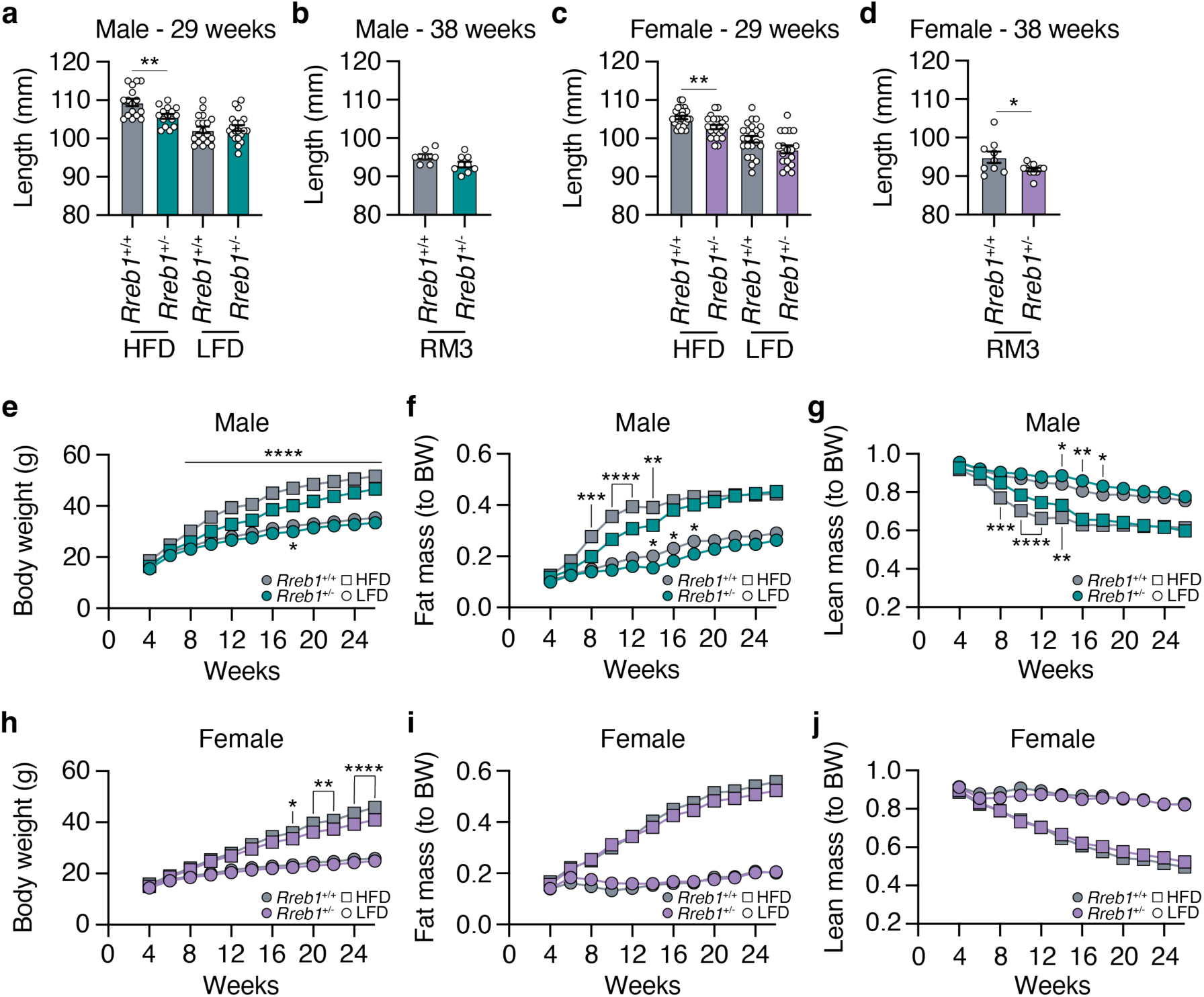
Reduced length, body weight, and fat mass in *Rreb1* heterozygous knockout mice. **a** Length in mm of wildtype (grey) and *Rreb1* heterozygous knockout (green) male mice at 29 weeks on HFD and LFD. *n* = 15-21. **b** Length in mm of wildtype (grey) and *Rreb1* heterozygous knockout (green) male mice at 38 weeks on RM3 diet. *n* = 7-9. **c** Length in mm of wildtype (grey) and *Rreb1* heterozygous knockout (purple) female mice at 29 weeks on HFD and LFD. *n* = 18-24. **d** Length in mm of wildtype (grey) and *Rreb1* heterozygous knockout (purple) female mice at 38 weeks on RM3 diet. *n* = 9-11. **e-g** Biweekly measurements of (**e**) body weight in grams (g), (**f**) fat mass normalized to body weight (BW), and (**g**) lean mass normalized to body weight (BW) of wildtype (grey) and *Rreb1* heterozygous knockout (green) male mice on HFD and LFD. *n* = 15-21. **h-j** Biweekly measurements of (**h**) body weight in grams (g), (**i**) fat mass normalized to body weight (BW), and (**j**) lean mass normalized to body weight (BW) of wildtype (grey) and *Rreb1* heterozygous knockout (purple) female mice on HFD and LFD. *n* = 18-24. Data are presented as mean ± s.e.m. Statistical analyses were performed by (**a-d**) unpaired t test (**d**, with Welch correction) or (**e-j**) two-way ANOVA with Tukey’s multiple comparisons test. \**p*<0.05, \*\**p*<0.01, \*\*\**p*<0.001, and \*\*\*\**p*<0.0001.

In addition to the reduction in body length, *Rreb1^+/-^*male mice had reduced body weight by 8 weeks of age on HFD (Fig. 1e). Biweekly EchoMRI was used to determine if the changes in body weight were due to alterations in body composition. Male *Rreb1*^+/-^ mice had decreased fat mass (Fig. 1f) and increased lean mass (Fig. 1g) when normalized to body weight from 8 to 14 weeks of age before converging with colony-mate controls at 16 weeks. While there was little change in body weight on LFD (Fig. 1e), male *Rreb1^+/-^* mice also weighed less on the RM3 diet (Extended Data Fig. 1a,b), had decreased fat mass (Extended Data Fig. 1c,d), and increased lean mass (Extended Data Fig. 1e,f). Female *Rreb1^+/-^* mice had reduced body weight on HFD at 18 weeks of age (Fig. 1h) and at 13 weeks of age on RM3 (Extended Data Fig. 1g,h), but had no significant differences in fat mass (Fig. 1i) or lean mass (Fig. 1j) on HFD, LFD, or on RM3 diet (Extended Data Fig. 1i,j,k,l). Overall, *Rreb1* heterozygosity, depending on diet and sex, resulted in mice that were smaller (shorter and lighter), with reduced relative fat mass and increased lean mass.

### Global *Rreb1* heterozygous knockout mice have altered white adipose tissue depot size

To determine if the decrease in overall fat mass changed the size of various visceral and subcutaneous fat depots, we measured fat depot size in male and female *Rreb1^+/-^* mice. On HFD, male *Rreb1* heterozygous mice had increased gonadal (Fig. 2a) and mesenteric visceral fat pads (Fig. 2b) but had no significant differences in perirenal (Fig. 2c) or subcutaneous inguinal fat pads (Fig. 2d) compared to wildtype control mice. Conversely, on LFD and RM3 diet male *Rreb1^+/-^* mice had decreased gonadal (Fig. 2a; Extended Data Fig. 2a) and on RM3 diet had decreased mesenteric (Extended Data Fig. 2a) visceral fat pads, which is consistent with the overall decrease in fat mass measured by EchoMRI. Female *Rreb1^+/-^* knockout mice also had decreased gonadal (Fig. 2e) and mesenteric (Fig. 2f) fat depot size on HFD, and no changes in white adipose depot size on LFD (Fig. 2e-h) or on RM3 diet (Extended Data Fig. 2b). As RREB1 is a transcription factor, we isolated brown, gonadal, and inguinal adipose tissues of *Rreb1*^+/+^ and *Rreb1*^+/-^ mice on RM3 diet at 26 weeks of age and performed bulk RNA-seq. Differential expression analysis identified 70 genes that were significantly upregulated (padj < 0.05 and log2FC > 1.5) and only two that were downregulated (padj < 0.05 and log2FC < -1.5) (Extended Data Table 1) in adipose tissue isolated from adult mice. Gene enrichment analysis found that the differentially expressed genes were enriched for GO terms relating to muscle, such as ‘sarcomere organization’, ‘myofibril assembly’, and ‘muscle cell development’ (Extended Data Table 2). Given the differences in fat mass we next assessed whether *Rreb1^+/-^* mice had changes in plasma adiponectin levels. Adiponectin levels were unchanged at 12 weeks (Fig. 2i) and significantly increased at 22 weeks (Fig. 2j) in *Rreb1^+/-^* male mice on HFD, consistent with the increased gonadal and mesenteric fat mass. There were no changes in adiponectin level in *Rreb1*^+/-^ male mice on LFD (Fig. 2i,j) or on RM3 diet (Extended Data Fig. 2c,d) and no differences in adiponectin levels in female *Rreb1^+/-^* mice (Fig. 2k,l and Extended Data Fig. 2e,f), despite reductions in fat depot size.

**Fig. 2.**
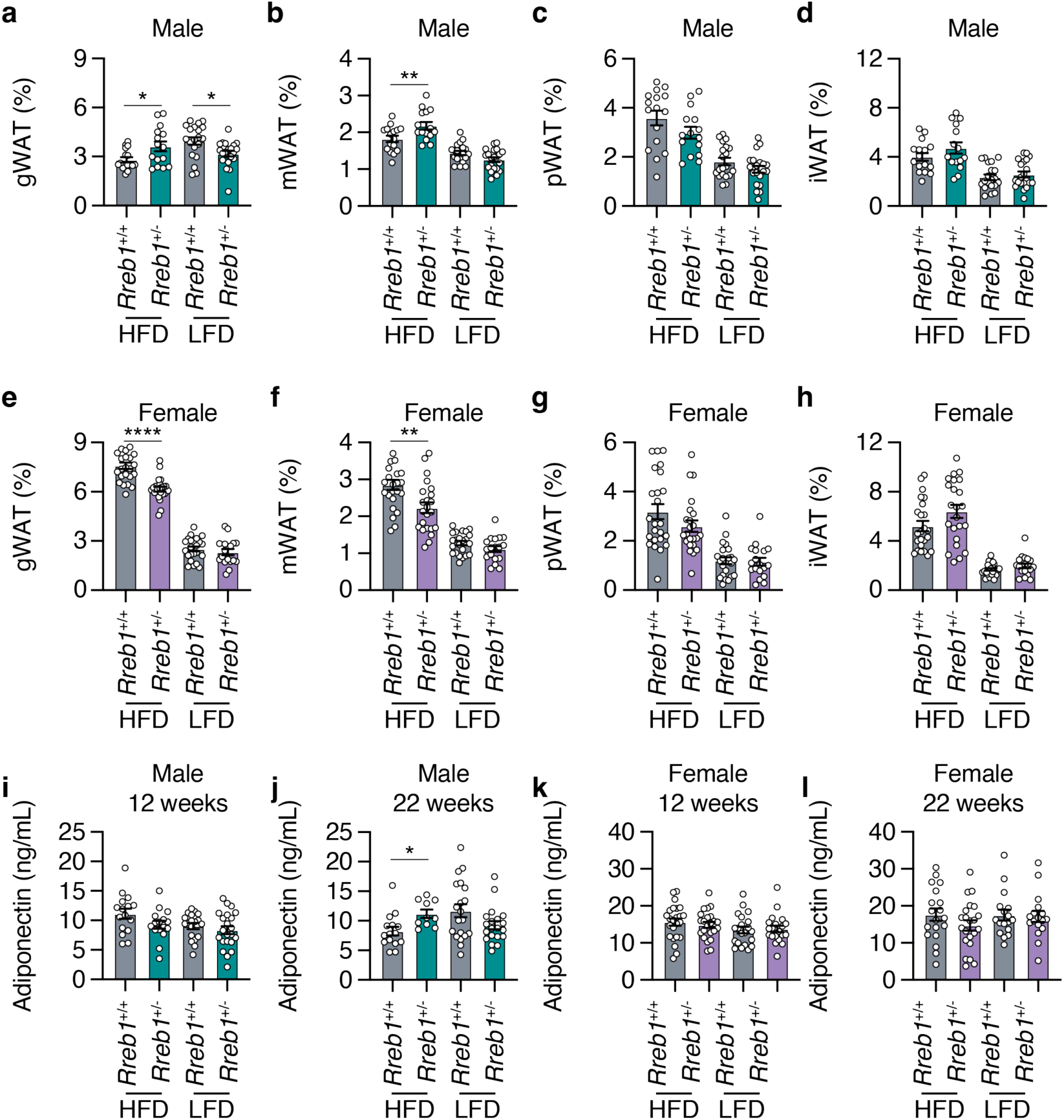
*Rreb1* heterozygous knockout mice have differences in depot size of white adipose tissue. **a-d** Comparisons of visceral gonadal (**a,** gWAT), mesenteric (**b,** mWAT), perirenal (**c,** pWAT), and subcutaneous inguinal (**d,** iWAT) white adipose tissue weight (% normalized to body weight) at 26 weeks in wildtype (grey) and *Rreb1* heterozygous knockout (green) male mice on HFD and LFD. *n* = 15-21. **e-h** Comparisons of visceral gonadal (**e,** gWAT), mesenteric (**f,** mWAT), perirenal (**g,** pWAT), and subcutaneous inguinal (**h,** iWAT) white adipose tissue weight (% normalized to body weight) at 26 weeks in control (grey) and *Rreb1* heterozygous knockout (purple) female mice on HFD and LFD. *n* = 18-24. **i,j** Plasma adiponectin (ng/mL) after an overnight fast at (**i**) 12 weeks and (**j**) 22 weeks for male wildtype (grey) and *Rreb1* heterozygous knockout (green) mice on HFD and LFD. *n* = 15-22. **k,l** Plasma adiponectin (ng/mL) after an overnight fast at (**k**) 12 weeks and (**l**) 22 weeks for female wildtype (grey) and *Rreb1* heterozygous knockout (purple) mice on HFD and LFD. *n* = 17-24. Data are presented as mean ± s.e.m. Statistical analyses were performed using a one-way ANOVA with Sidak’s multiple comparison test or (**f,g,h,j**) Brown-Forsythe and Welch ANOVA tests with Dunnett’s T3 multiple comparisons test \**p*<0.05, \*\**p*<0.01, and \*\*\*\**p*<0.0001.

To understand the reduction of adipose tissue depot size in *Rreb1*^+/-^ mice, we next assessed the distribution of adipocyte size and cell number by histological analysis of white adipose tissues at 29 weeks of age. In the gonadal depot, male *Rreb1*^+/-^ mice on HFD had significantly fewer adipocytes of smaller size (area <4500 µm^2^) and more adipocytes of larger size (area >5750 µm^2^) (Fig. 3a), consistent with the increased gonadal depot size (Fig. 2a). In females, gonadal adipocyte size was unchanged for *Rreb1*^+/-^ mice on HFD (Fig. 3b), despite a smaller overall gonadal tissue size (Fig. 2e). For subcutaneous adipocytes, there was no significant difference in adipocyte size in male *Rreb1*^+/-^ mice on HFD (Fig. 3c). However, female *Rreb1*^+/-^ mice had significantly more subcutaneous adipocytes of smaller size (area <4500 µm^2^) and fewer adipocytes of larger size (>5250 µm^2^) compared to wildtype mice on HFD (Fig. 3d). Similarly to females, male *Rreb1*^+/-^ mice on a RM3 diet at 38 weeks of age had a significant increase in the number of small adipocytes and fewer large adipocytes in gonadal (Extended Data Fig. 3a), perirenal (Extended Data Fig. 3b), and subcutaneous adipose tissue (Extended Data Fig. 3d) compared to wildtype control mice. Finally, there were no differences in visceral gonadal (Fig. 3e,f), mesenteric (Extended Data Fig. 3c) or subcutaneous (Fig. 3g,h) adipocyte size for *Rreb1*^+/-^ mice on LFD or RM3 diet. Taken together, these results suggest that haploinsufficiency of *Rreb1* leads to sex- and diet-specific decreases in fat mass, white adipose depot size, adipocyte size, and adipocyte area.

**Fig. 3.**
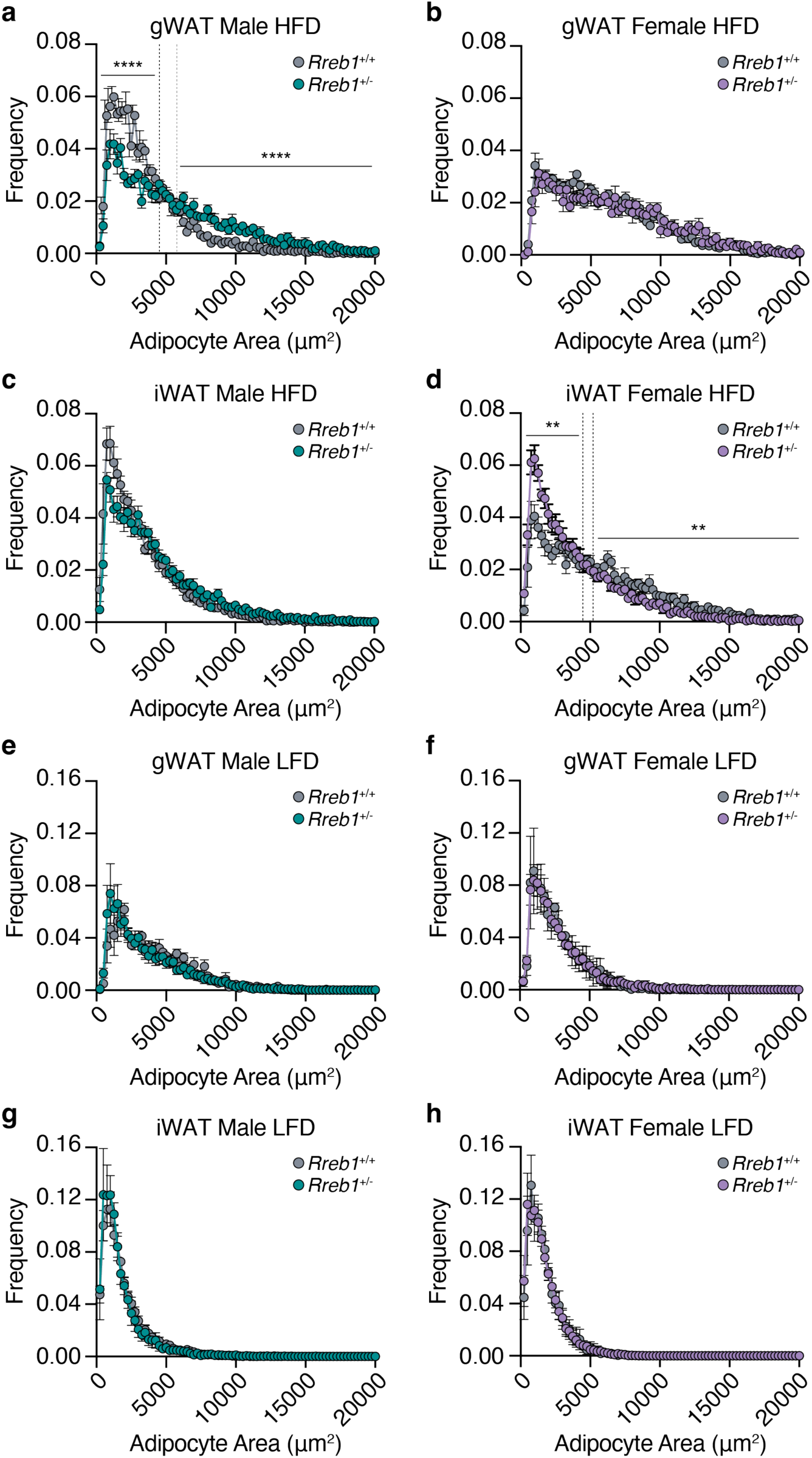
Sex- and tissue-specific differences in adipocyte size of white adipose tissue in *Rreb1* heterozygous knockout mice. **a-d** Quantification of adipocyte area (µm^2^) within (**a,b**) visceral gonadal (gWAT, both sexes: *Rreb1*^+/+^ *n*= 5, *Rreb1*^+/-^ *n* = 5 mice) and (**c,d**) subcutaneous inguinal (iWAT, both sexes: *Rreb1*^+/+^ *n* = 6, *Rreb1*^+/-^ *n* = 6 mice) fat depots from (**a,c**) male and (**b,d**) female *Rreb1*^+/+^ and *Rreb1*^+/-^ mice fed a HFD at 29 weeks of age. Dotted line denotes the interval where the dataset converges. **e-h** Quantification of adipocyte area (µm^2^) within (**e,f**) visceral gonadal (gWAT, both sexes: *Rreb1*^+/+^ *n* = 3, *Rreb1*^+/-^ *n* = 6 mice) and (**g,h**) subcutaneous inguinal (iWAT, male: *Rreb1*^+/+^ *n* = 6, *Rreb1*^+/-^ *n* = 5; female: *Rreb1*^+/+^ *n*= 6, *Rreb1*^+/-^ *n* = 6 mice) fat depots from (**e,g**) male and (**f,h**) female *Rreb1*^+/+^ and *Rreb1*^+/-^ mice fed a LFD at 29 weeks of age. Data are presented as mean adipocyte frequency within each 250 µm^2^ sized bin ± s.e.m. Statistical analyses were performed on the area under the curve for adipocytes in (**a**) <4500 and >5750 µm^2^ or (**d**) <4500 and >5250 µm^2^ adipocyte size ranges by (**a,d,e,f,h**) unpaired t test or (**b,c,g**) Mann-Whitney two tailed test, depending on the normality of the data distribution. \*\**p*<0.01 and \*\*\*\**p*<0.0001.

### Global *Rreb1* knockout mice have increased lipid accumulation in the liver

To determine whether the reduced adipose tissue size was due to fat being stored ectopically in the liver, we next performed histological analysis of liver tissue in *Rreb1*^+/-^ male and female mice. *Rreb1*^+/-^ male and female mice had qualitatively increased lipid droplet accumulation in the liver compared to wildtype controls (Extended Data Fig. 4). To assess the impact of ectopic fat on liver function in *Rreb1*^+/-^ mice, serum alkaline phosphatase (ALP), alanine aminotransferase (ALT), and aspartate aminotransferase (AST) levels were measured. There were no differences in liver enzymes between male *Rreb1^+/-^*and control mice (Fig. 4a,b,c). However, female *Rreb1^+/-^* mice had significantly increased ALP, ALT, and AST levels (Fig. 4d,e,f). Consistent with the liver enzyme measurements, male *Rreb1^+/-^*mice had reduced liver weight (Fig. 4g), while female *Rreb1^+/-^* mice had increased liver weight compared to littermate controls (Fig. 4h). These data suggest that *Rreb1* heterozygosity leads to ectopic lipid accumulation in the liver.

**Fig. 4.**
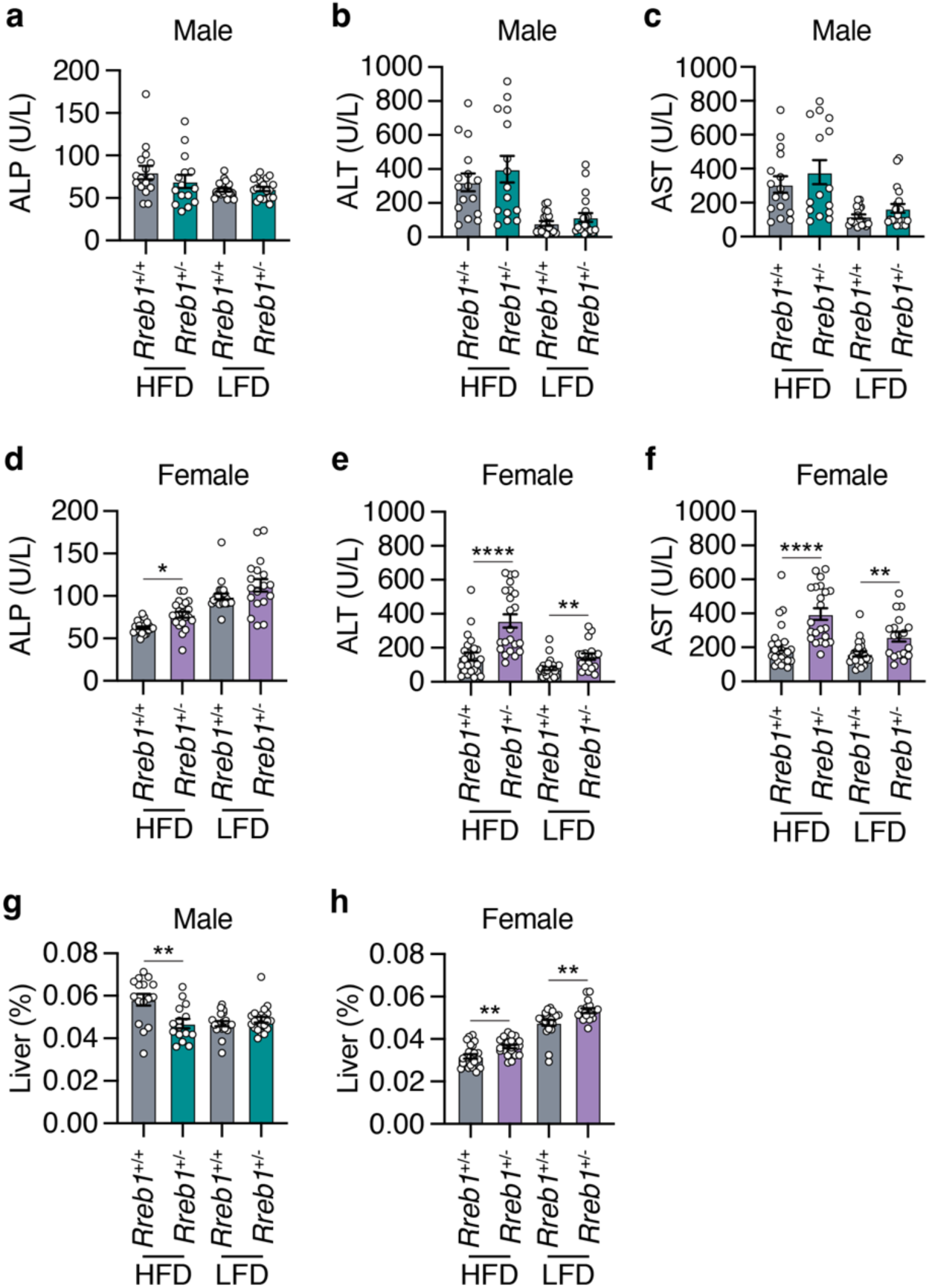
Global *Rreb1* heterozygous knockout mice have ectopic fat in the liver. **a,b,c** Plasma (**a**) alkaline phosphatase (ALP; U/L), (**b**) alanine aminotransferase (ALT; U/L), and (**c**) aspartate aminotransferase (AST; U/L) levels in *Rreb1*^+/+^ and *Rreb1*^+/-^ male mice at 29 weeks on HFD and LFD. *n* = 15-20. **d,e,f** Plasma (**d**) alkaline phosphatase (ALP; U/L), (**e**) alanine aminotransferase (ALT; U/L), and (**f**) aspartate aminotransferase (AST; U/L) levels in *Rreb1*^+/+^ and *Rreb1*^+/-^ female mice at 29 weeks on HFD and LFD. *n* = 14-23. **g,h** Liver weight (adjusted for body weight, %) of *Rreb1*^+/+^ and *Rreb1*^+/-^ on HFD and LFD at 29 weeks of age in (**g**) male (*n* = 15-21) and (**h**) female (*n* = 17-23) mice. Data are presented as mean ± s.e.m. Statistical analysis performed by (**a**,**g**) Brown-Forsythe and Welch ANOVA tests with Dunnett’s T3 multiple comparisons test, (**b**,**c**,**e**,**f**) ordinary one-way ANOVA with Sidak’s multiple comparison tests, and (**d**) ANOVA Kruskal-Wallis test. (**a**,**b**,**e,f**) data were transformed Y=log(Y) before statistical analysis. * *p*<0.05, ** *p*<0.01, and **** *p*<0.0001.

### Global heterozygous *Rreb1* knockout mice show reduced food intake and energy expenditure

To test whether the reduced weight and fat mass in *Rreb1*^+/-^ mice is due to changes in food intake and energy expenditure, we measured weekly food intake from 6 to 24 weeks on HFD. Both male (Fig. 5a) and female *Rreb1*^+/-^ mice (Fig. 5b) had reduced food intake compared to colony-mate controls (AUC: males, *p* = 0.013; females, *p* = 0.0126). After 24 weeks, energy expenditure was assessed by indirect calorimetry. While male *Rreb1*^+/-^ mice had no change in heat production (Fig. 5c), female *Rreb1*^+/-^ mice had significantly decreased heat production (Fig. 5d), consistent with reduced energy expenditure. There were no changes in the percentage of brown adipose tissue (BAT) in either male (Fig. 5e) or female (Fig. 5f) *Rreb1*^+/-^ mice, suggesting no thermogenic differences due to *Rreb1* haploinsufficiency. The same cohorts of mice were also assessed for body weight and composition during the food intake phenotyping period. Interestingly, *Rreb1*^+/-^ male mice no longer showed statistically significant differences in body weight and fat mass compared to the controls (Extended Data Fig. 5a,c), potentially due to the effects of housing two male mice together. The body weight of *Rreb1*^+/-^ female mice was significantly decreased (Extended Data Fig. 5b), consistent with previous observations, but additionally both fat and lean mass were also decreased in these mice (Extended Data Fig. 5d,f). Consistent with decreases in body weight and fat content, serum leptin was unchanged in male *Rreb1*^+/-^ mice (Fig. 5g), but significantly decreased in female *Rreb1*^+/-^ mice (Fig. 5h). Therefore, *Rreb1*^+/-^ mice have reduced adiposity, decreased food intake, and decreased circulating leptin levels.

**Fig. 5.**
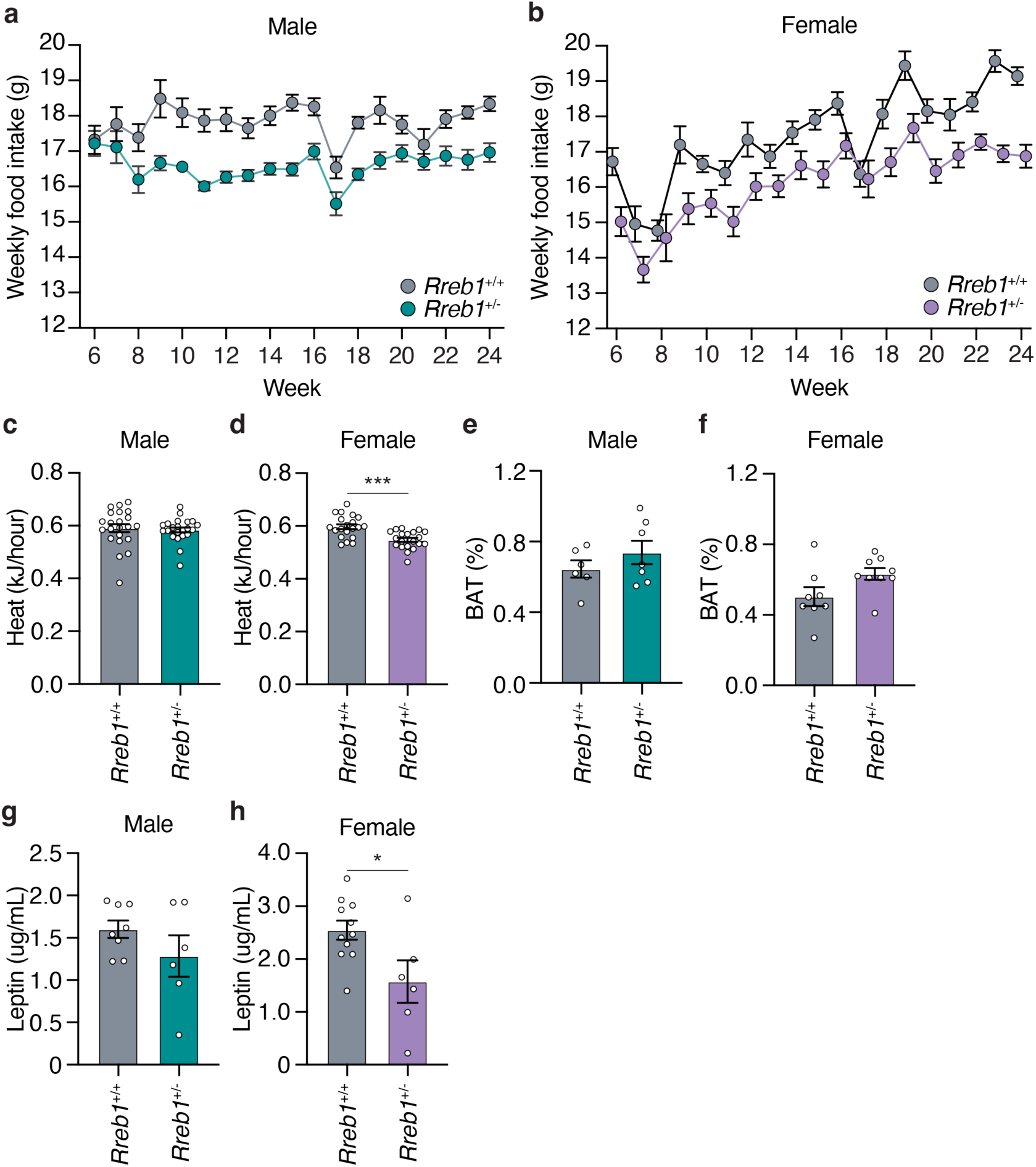
Reduced food intake and energy expenditure in *Rreb1* heterozygous knockout mice. **a,b** Comparisons of weekly food intake (g) measured from 6 to 24 weeks for *Rreb1*^+/+^ and *Rreb1*^+/-^ (**a**) male and (**b**) female mice on HFD. (**a**) *Rreb1*^+/+^ and *Rreb1*^+/-^ *n* = 11 cages, each containing 2 animals. (**b**) *Rreb1*^+/+^ *n* = 9 and *Rreb1*^+/-^ *n* = 11 cages, each containing 2 animals. **c,d** Heat production of *Rreb1*^+/+^ and *Rreb1*^+/-^ (**c**) male and (**d**) female mice over a 24-hour period. (**c**) *n* = 22, (**d**) *n* = 20 mice. **e,f** Comparisons of brown adipose tissue (BAT) weight of *Rreb1*^+/-^ (**e**) male and (**f**) female mice with wildtype littermate control mice at 38 weeks on RM3 diet. (**e**) *n* = 6-7 mice, (**f**) *n =* 8-9 mice. **g,h** Circulating leptin levels in *Rreb1*^+/-^ (**g**) male and (**h**) female mice with wildtype littermate control mice. (**g**) *n* = 6-8, (**h**) *n* = 6-11 mice. Data are presented as mean ± s.e.m. (**a**,**b**) Area under the curve was calculated for statistical analysis (males: 6-24 weeks and females: 6-23 weeks). Statistical analyses were performed by two-tailed unpaired t test or (**c**) a Mann Whitney test. \**p*<0.05, \*\*\**p*<0.001.

### Global *Rreb1* heterozygous knockout mice exhibit improved insulin sensitivity

To determine the impact of *Rreb1* haploinsufficiency on glucose homeostasis, we next performed intraperitoneal glucose tolerance tests (IPGTT) at 12 and 22 weeks of age. Neither male (Extended Data Fig. 6a,b) nor female (Extended Data Fig. 6c,d) *Rreb1*^+/-^ mice had differences in glucose tolerance. Insulin sensitivity was measured using an intraperitoneal insulin sensitivity test (IPIST) at 16 and 26 weeks of age. Male *Rreb1*^+/-^ mice fed a HFD had improved insulin sensitivity at 16 weeks of age (Fig. 6a), but had no significant differences in insulin sensitivity at 26 weeks of age (Fig. 6b). However, male *Rreb1*^+/-^ mice had reduced insulin sensitivity at 26 weeks of age when fed a LFD (Fig. 6b). No differences in insulin sensitivity were detected for *Rreb1*^+/-^ female mice on HFD or LFD at either 16 (Fig. 6c) and 26 weeks of age (Fig. 6d), which may result from female mice being protected against HFD-induced metabolic syndrome^25^. In line with the early improvement in insulin sensitivity, fasting insulin levels were decreased in male *Rreb1*^+/-^ mice at 12 weeks (Fig. 6e) and 22 weeks (Fig. 6f) on HFD. Female *Rreb1*^+/-^ mice had no change in fasting insulin at 12 or 22 weeks of age on HFD or LFD (Fig. 6g,h). As we have previously shown that loss of *Rreb1* negatively impacts beta cell function^21^, we measured insulin secretion from mouse islets *ex vivo*. Insulin secretion in response to high glucose from *Rreb1*^+/-^ female mouse islets was significantly reduced (Extended Data Fig. 6e).

**Fig. 6.**
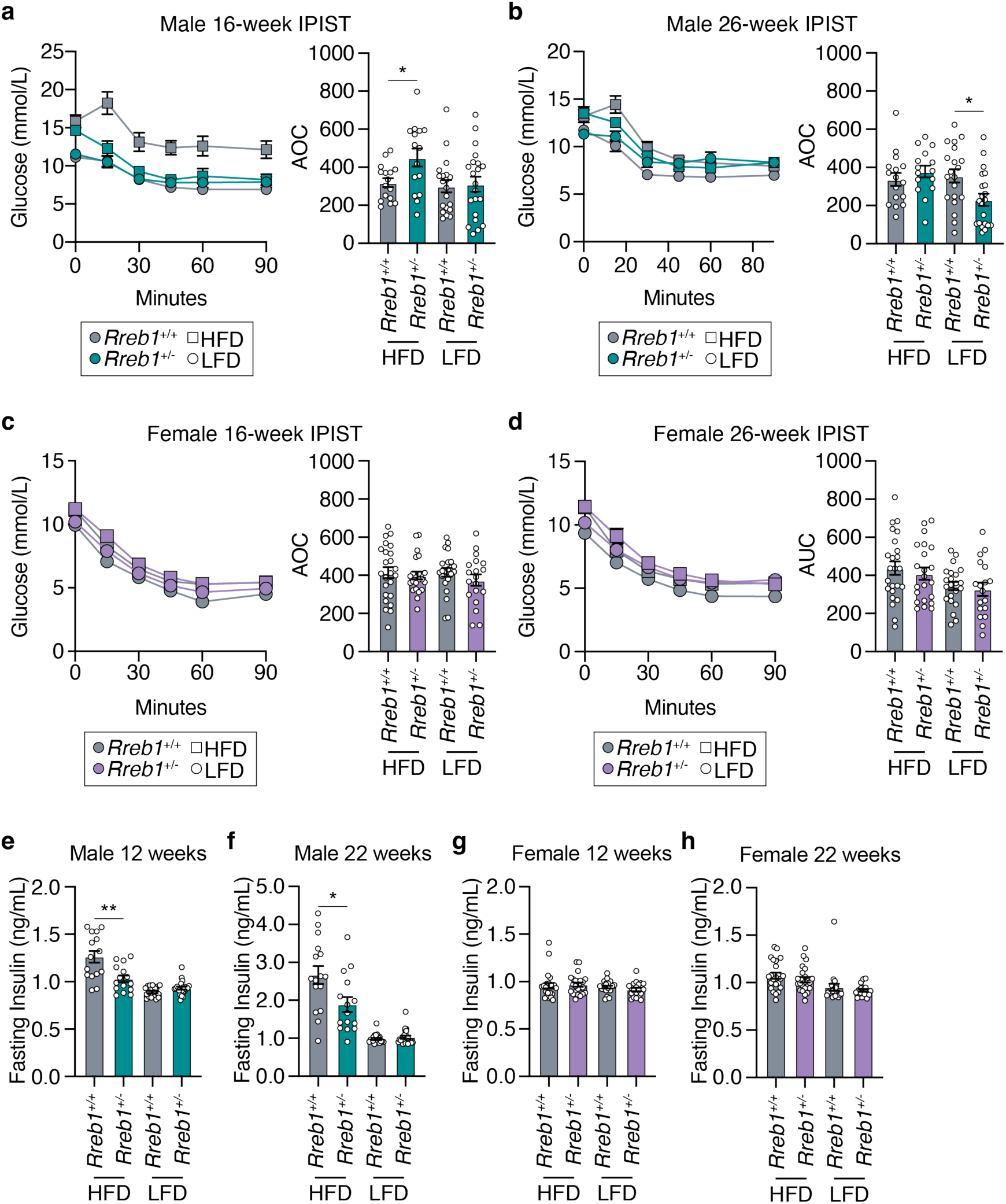
Male *Rreb1*^+/-^ mice have differences in insulin sensitivity and fasting insulin levels. **a,b** IPIST on *Rreb1*^+/+^ (grey) and *Rreb1*^+/-^ (green) male mice aged (**a**) 16 weeks and (**b**) 26 weeks on HFD and LFD *n* = 15-21. **c,d** IPIST on *Rreb1*^+/+^ (grey) and *Rreb1*^+/-^ (purple) female mice aged (**c**) 16 and (**d**) 26 weeks of age on HFD and LFD. *n* = 19-24. **e-h** Plasma insulin after an overnight fast at (**e,g**) 12 and (**f,h**) 22 weeks for (**e,f**) male and (**g,h**) female *Rreb1*^+/+^ and *Rreb1*^+/-^ mice on HFD and LFD. *n* = 15-24. Data are shown as mean ± s.e.m. Area of curve (AOC) was calculated and statistical analyses performed between genotypes using (**a,b,c,d**) one way ANOVA with Sidak’s multiple comparisons test or (**e,f,g**) Brown-Forsythe and Welch ANOVA tests with Dunnett’s T3 multiple comparisons test. Where necessary to normalize data distribution before analysis, data was transformed by (**e,h**) Y=1/Y or (**f,g**) Y=log(Y). \**p*<0.05, \*\**p*<0.01.

To determine if loss of *Rreb1* changed glucose uptake in adipose tissue, we performed radiolabeled tracing using [^3^H]-2-deoxy-glucose in wildtype and *Rreb1*^+/-^ mice on a standard chow diet (18% fat, 24% protein, and 58% carbohydrates) between age 34 and 37 weeks of age. Under basal conditions, male *Rreb1*^+/-^ mice had increased glucose uptake in perirenal white adipose tissue and decreased glucose uptake in brown adipose tissue (Fig. 7a). Moreover, when male *Rreb1*^+/-^ mice were stimulated with insulin, there was a significant increase in glucose uptake across all white adipose tissues (Fig. 7b). Female *Rreb1*^+/-^ had no change in glucose uptake under basal (Fig. 7c) or insulin stimulated conditions (Fig. 7d), consistent with the *in vivo* insulin sensitivity tests (Fig. 6c,d). Taken together, these results suggest that depending on sex, diet, and age, global *Rreb1* haploinsufficiency decreases insulin secretion, reduces fasting insulin, and results in smaller, more insulin sensitive adipocytes.

**Fig. 7.**
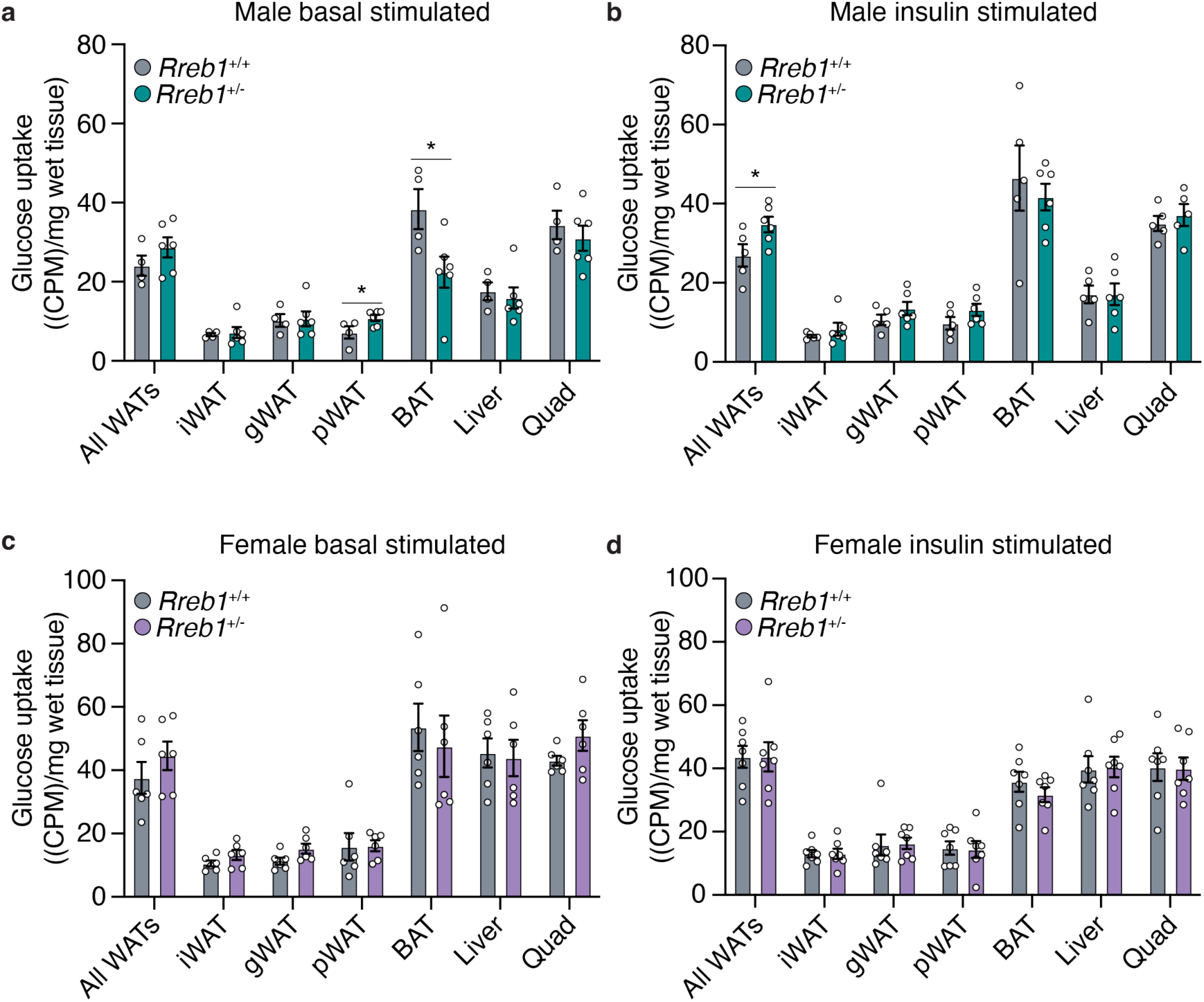
Improved insulin-stimulated glucose uptake in male *Rreb1^+/-^* mice. **a,b** Basal (**a**) and insulin-stimulated (**b**) glucose uptake ((CPM)/mg of wet tissue weight) in all white adipose tissues (WAT), inguinal (iWAT), gonadal (gWAT), perirenal (pWAT), brown adipose tissue (BAT), liver and quadricep in male wildtype (grey) and *Rreb1^+/-^* (green) mice. **c,d** Basal (**c**) and insulin-stimulated (**d**) glucose uptake ((CPM)/mg of wet tissue weight) in all white adipose tissues (WAT), inguinal (iWAT), gonadal (gWAT), perirenal (pWAT), brown adipose tissue (BAT), liver and quadricep in female wildtype (grey) and *Rreb1^+/-^* (purple) mice. Data are shown as mean ± s.e.m. Statistical analyses were performed between genotypes using an unpaired t-test. *n* = 4-7. \**p*<0.05.

### *Rreb1* perturbation impacts adipogenesis *in vitro*

To understand how loss of *Rreb1* reduces adipocyte size, we isolated cells from the mouse stromal vascular fraction (SVF) of *Rreb1*^+/+^ and *Rreb1*^+/-^ mice and differentiated them *in vitro* for 48 hours. There was a significant reduction in the formation of adipocytes from SVF cells isolated from male *Rreb1*^+/-^ mice compared to wildtype control (Fig. 8a). In addition, there was a significant increase in the number of BrdU+ S-phase cells immediately following induction of differentiation (Fig. 8b), suggesting that in the absence of Rreb1 fewer pre-adipocytes exit the cell cycle and commit to differentiation^26^.

**Fig. 8.**
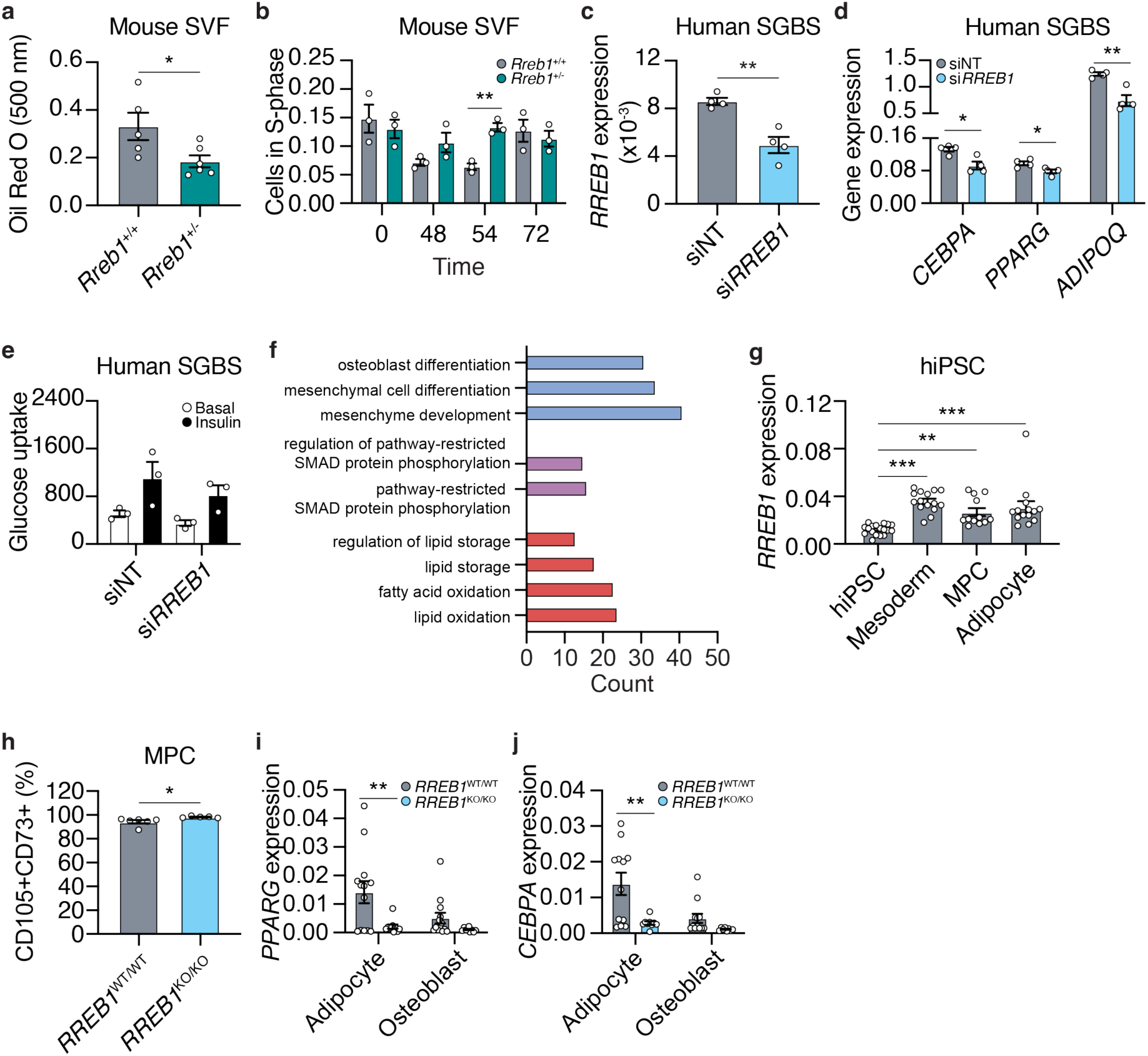
Loss of *Rreb1* decreases adipocyte formation and expression of pro-adipogenic genes. **a** Quantification of adipocyte formation using Oil Red O staining (absorbance at 500 nm) in stromal vascular fraction (SVF) cells from wildtype (*Rreb1*^+/+^; grey) and knockout (*Rreb1*^+/-^; green) mice. *n* = 5-6. **b** Quantification of BrdU+ S-phase cells during *in vitro* differentiation of SVF cells from *Rreb1*^+/+^ (grey) and *Rreb1*^+/-^ (green) male mice. *n* = 3. **c** *RREB1* expression 48 hours following transient transfection of siRNAs against *RREB1* (si*RREB1*) or non-targeting control (siNT) siRNAs in SGBS cells. *n* = 4. **d** Gene expression analysis of *CEBPA*, *PPARG*, and *ADIPOQ* at day 5 of *in vitro* differentiation of SGBS cells to adipocytes. SGBS were treated with siNT and si*RREB1* 48 hours before differentiation. *n* = 4. **e** Glucose uptake of SGBS cells at day 12 of *in vitro* differentiation. SGBS were treated with siNT and si*RREB1* 48 hours before differentiation. *n* = 3. **f** Gene enrichment analysis of *RREB1* knockdown SGBS cells at day 12. The number of differentially expressed genes (count) in a subset of gene ontologies relating to mesenchymal stem cell differentiation (blue), SMAD pathway (purple), and lipids (red). **g** *RREB1* transcript expression normalized to *PPIA* in human induced pluripotent stem cell (hiPSC), mesoderm, mesenchymal progenitor cells (MPC), and adipocytes. **h** Flow cytometry analysis of CD105 and CD73 co-expression in wildtype (*RREB1*^WT/WT^; grey) and *RREB1* knockout (*RREB1*^KO/KO^; blue) hiPSC-derived mesenchymal progenitor cells (MPC). **i,j** hiPSC wildtype (*RREB1*^WT/WT^; grey) and *RREB1* knockout (*RREB1*^KO/KO^; blue) cells were differentiated to adipocyte and osteoblast lineages. The expression of adipocyte genes (**i**) *PPARG* and (**j**) *CEBPA* were measured by qPCR and normalized to *PPIA*. Data are presented as mean ± s.e.m. Statistical analyses were performed by unpaired t test or one-way ANOVA. \**p*<0.05, \*\**p*<0.01, \*\*\**p*<0.001.

To model human adipogenesis we used the Simpson-Golabi-Behmel syndrome (SGBS) human pre-adipocyte cell model^27^. *RREB1* transcript expression increased over the first four days of adipocyte induction and continued to be expressed in maturing adipocytes (Day 8 and 12) (Extended Data Fig. 7a,b), coinciding with the upregulation of pro-adipogenic genes (Extended Data Fig. 7c,d,e; Extended Data Table 3). Supporting our data, *RREB1* transcript was also found to be significantly upregulated in scRNA-seq data of adipocytes derived from SGBS cells^28^. Additionally, *RREB1* is more highly expressed in human adipocytes than in progenitor cells^29,30^, consistent with a potential role for RREB1 in human adipogenesis.

To determine if RREB1 is required for adipogenesis, *RREB1* expression was transiently knocked down using siRNAs two days before differentiation; gene expression analysis confirmed a 57% reduction in *RREB1* transcript at day 0 (Fig. 8c). After five days of differentiation, loss of *RREB1* significantly reduced expression of adipogenic genes *CEBPA*, *PPARG*, and *ADIPOQ* (Fig. 8d), suggesting RREB1 is required for human adipogenic differentiation. Interestingly, despite defects in adipogenic gene expression, there were no differences in insulin-stimulated glucose uptake (Fig. 8e), even when normalized for total protein content (Extended Data Fig. 7f). Transcriptomic analysis of siNT control and si*RREB1* SGBS cells after five days of differentiation identified 170 upregulated (log2FC >1.5, padj < 0.05) and 73 downregulated (log2FC < -1.5, padj < 0.05) genes following *RREB1* knockdown (Extended Data Table 4). Gene enrichment analysis revealed a significant enrichment of genes involved in ‘regulation of pathway-restricted SMAD protein phosphorylation’ and ‘pathway-restricted SMAD protein phosphorylation’ (Fig. 8f; Extended Data Table 5), which is consistent with the known interaction of RREB1 and SMAD proteins to activate gene expression during epithelial-to-mesenchymal transition^15^. There were significant enrichments for several GO terms relating to lipids, including ‘regulation of lipid storage’, ‘lipid storage’, ‘fatty acid oxidation’, and ‘lipid oxidation’ (Fig. 8f; Extended Data Table 5). Interestingly, the GO terms ‘osteoblast differentiation’, ‘mesenchymal cell differentiation’, and ‘mesenchyme development’ were significantly enriched (Fig. 8f; Extended Data Table 5), suggesting that loss of *RREB1* results in activation of osteogenic genes.

To understand if RREB1 is required for mesenchymal progenitor cell differentiation to fat, we used a *RREB1* homozygous null human induced pluripotent stem cell (hiPSC) line^21^. Isogenic wildtype control (*RREB1*^WT/WT^) and knockout (*RREB1*^KO/KO^) hiPSCs were differentiated to mesoderm, mesenchymal progenitor cells (MPC), and adipocytes using established protocols^31^ and commercially available kits. *RREB1* transcript is detected throughout all stages of the differentiation and peaks in mesodermal cells (Fig. 8g), consistent with the known role of RREB1 in early development and gastrulation^22^. *RREB1* transcript was significantly reduced at the mesoderm stage (Extended Data Fig. 7g). *RREB1* knockout significantly increased the expression of key mesodermal transcription factors *MIXL1* (Extended Data Fig. 7h), *TBXT* (Extended Data Fig. 7i), without changing *NCAM1* expression (Extended Data Fig. 7j). There was also a significant increase in the formation of CD105+CD73+ MPCs derived from *RREB1*^KO/KO^ hiPSCs compared to wildtype control (Fig, 8h). Despite improved formation of earlier progenitor populations, *RREB1*^KO/KO^ MPCs differentiated to both adipocyte and osteoblast lineages had reduced expression of adipocyte genes *PPARG* (Fig. 8i) and *CEBPA* (Fig. 8j). Together, these data suggest that in the absence of RREB1 MPCs, the common precursor to both adipocytes and osteoblasts, fail to robustly activate expression of adipogenic genes and instead activate the osteoblast lineage.

### *Rreb1* haploinsufficiency increases bone mineral density in mice

As the *RREB1* locus is also associated with bone mineral density (BMD)^7^ and our *in vitro* differentiation model supports changes in osteoblast formation, we measured BMD in the tibia of *Rreb1*^+/-^ mouse using dual-energy X-ray absorptiometry (DEXA). While there were no differences in BMD of male *Rreb1^+/-^*mice (Fig. 9a), female *Rreb1*^+/-^ mice on HFD and LFD at 29 weeks had significantly increased BMD (Fig. 9b). Further measurements of the tibial trabecular bone using microCT found no significant differences in trabecular thickness (Fig. 9c; Tb.Th) and separation (Fig. 9d; Tb.Sp) following *Rreb1* knockout. However, there were significant increases in both trabecular bone volume (Fig. 9e; BV/TV) and trabecular number (Fig. 9f; Tb.N) in female mice. Together, these data suggest that loss of *RREB1* influences skeletal traits and is directionally consistent with human genetic association data for the *RREB1* locus.

**Fig. 9.**
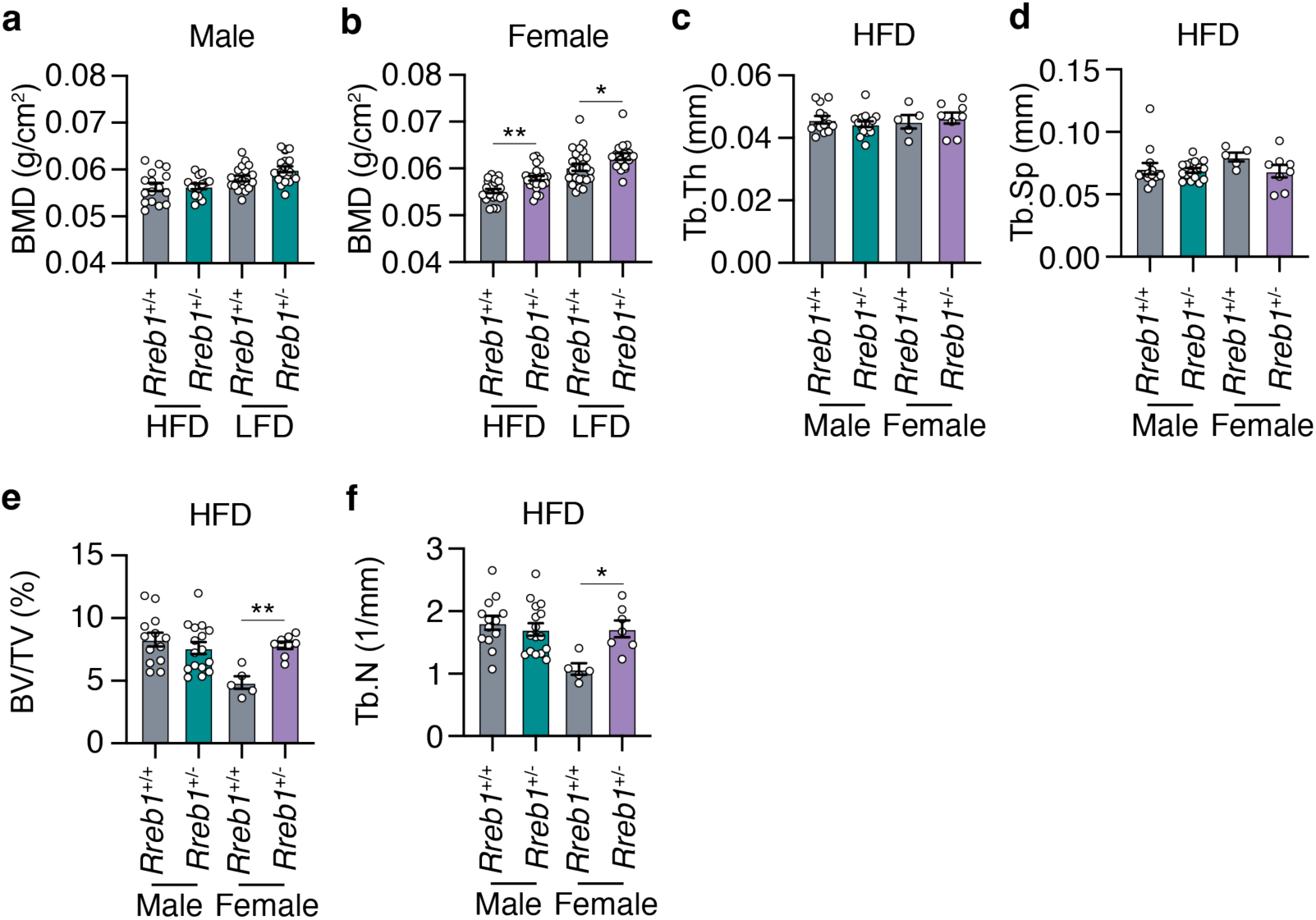
Increased bone mineral density in *Rreb1* heterozygous knockout mice. **a,b** Bone mineral density (BMD; g/cm^2^) on HFD and LFD of (**a**) male (*n* = 14-20) and (**b**) female (*n* = 17-24) mice at 29 weeks. **c,d,e,f** MicroCT analysis of male and female mice on HFD for (**c**) trabecular thickness (Tb.Th), (**d**) trabecular separation (Tb.Sp), (**e**) bone volume/tissue volume (BV/TV), and (**f**) trabecular number (Tb.N). *n* = 5-16. Data are presented as mean ± s.e.m. Statistical analyses were performed by one-way ANOVA with Šídák’s multiple comparisons test. *p<0.05 and **p<0.01.

### Human carriers of *RREB1* non-coding protective alleles have reduced adipocyte area

A consistent phenotype amongst our *in vitro* and *in vivo* models is a reduction in adipocyte size. To understand if human carriers of T2D-protective alleles have alterations in adipocyte size, we performed histological analysis of adipocyte area in donor subcutaneous and visceral adipose tissue^32^. For two of the T2D association signals (rs9505097 and rs9370984) at the *RREB1* locus, there were no significant differences in subcutaneous or visceral adipocyte area (Extended Data Fig. 8). For the rs112498319 variant, homozygous carriers of the T2D-protective allele had a significant reduction in subcutaneous adipocyte area in all individuals (Fig. 10a). When classifying individual donors by sex, significant reductions in subcutaneous adipocyte area for both male and female carriers of the protective A allele remained (Fig. 10a). While there was also a significant reduction in visceral adipocyte area in all individuals, the reduction was driven by a significant reduction in female carriers of the T2D-protective A allele (Fig. 10b). The credible set for the rs112498319 contains over 200 variants (Extended Data Fig. 9), one of which is located within a region that has long-range contact with the *RREB1* gene (Extended Data Fig. 9). This region overlaps with a putative enhancer that is activated during adipogenesis based on ATAC-seq data from SGBS cells as well as MED1 and H3K27ac ChIP-seq data and DNase I hypersensitive sites from BM-hMSCs (Extended Data Fig. 9). Taken together, these data support our mouse model where *Rreb1* heterozygous loss-of-function is protective against T2D by reducing adipocyte size in females, leading to more insulin sensitive adipocytes.

**Fig. 10.**
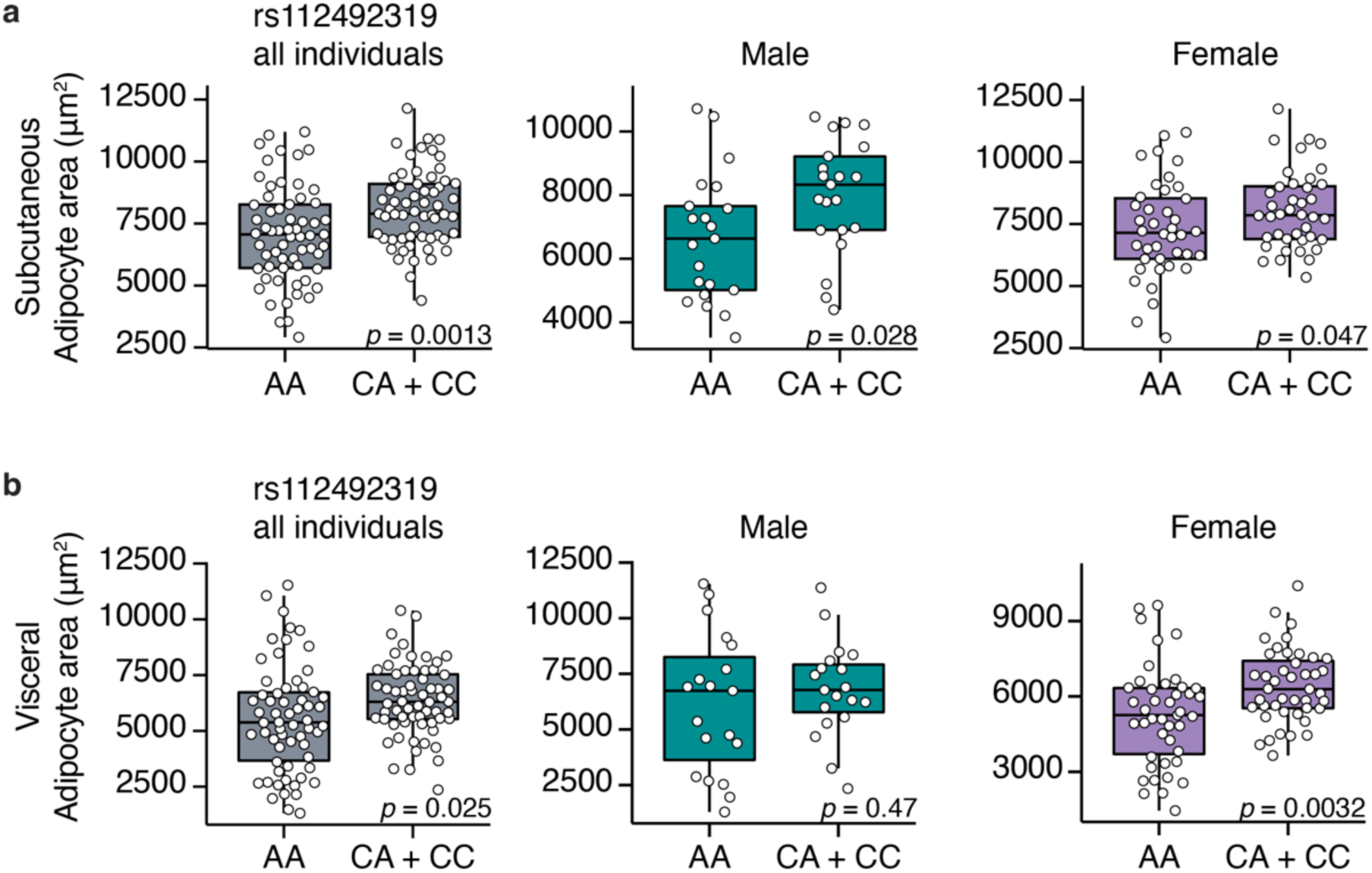
Changes in human adipocyte area in *RREB1* rs112492319 variant carriers. **a,b** Adipocyte area of (**a**) subcutaneous and (**b**) visceral fat from human donors carrying rs112492319 variants. Left graph: all individuals (pooled and matched); middle graph: males; right graph: females. Data are presented as mean ± s.e.m. Statistical analyses were performed by unpaired t-test. Individual *p* values are labelled in each graph.

## Discussion

There is compelling genetic evidence that variation at the *RREB1* locus influences T2D risk and other metabolic traits. Our previous study^21^ supports a role for pancreatic beta cells in mediating T2D risk at the *RREB1* locus; however, the relative contribution of other diabetes-relevant tissues and whether *RREB1* T2D-risk alleles result in a loss- or gain-of-function remains unknown. Using a global heterozygous *Rreb1* knockout mouse model, we now show that *Rreb1* haploinsufficiency reduces length, body weight, and fat mass (Fig. 11). *Rreb1*^+/-^ mice had diet- and sex-specific differences in adipose tissue; male *Rreb1*^+/-^ mice on HFD had increased depot size and larger adipocytes, whereas male mice on LFD and female mice on HFD had reduced depot size with smaller adipocytes in females (Fig. 11). Male mice showed significant improved insulin sensitivity on HFD when measured *in vivo* by IPIST at 16 weeks (although lost by 26 weeks and reversed on a LFD at that age), together with lower fasting insulin on HFD at both age points, and had improved glucose uptake in white adipose tissue in response to insulin *in vivo*. Mouse and human pre-adipocytes demonstrated defects in adipogenesis and were instead primed to form osteoblasts, consistent with the increase in BMD measured in female *Rreb1*^+/-^ mice. Finally, there were significant decreases in adipocyte size for male and female human carriers of the T2D-protective alleles, suggesting that Rreb1 loss-of-function protects against diabetes due to the generation of smaller, more insulin sensitive adipocytes.

**Fig. 11.**
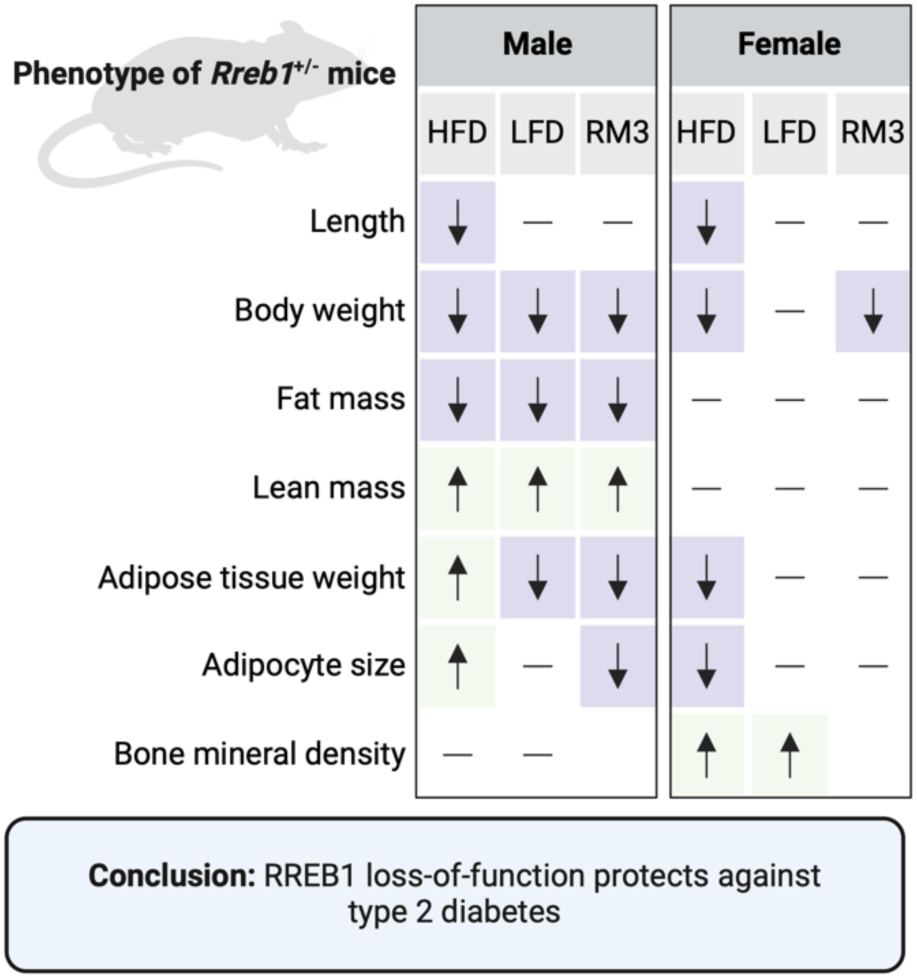
RREB1 loss-of-function protects against type 2 diabetes. Overview of significant differences in measured phenotypes in the global *Rreb1* heterozygous knockout male and female mice on HFD, LFD, and RM3 diet. Created with BioRender.com.

*Rreb1* haploinsufficiency in mice causes decreased weight gain on a HFD and diet-specific differences in visceral adipose tissue. Adipocyte tissue is expanded by either increasing the size (hypertrophy) or number (hyperplasia) of adipocytes. Adipocyte hypertrophy is associated with T2D^33,34^, dyslipidemia^35^, and cardiovascular disease^36^. Studies examining functional differences, such as gene expression^37,38^, adipokine secretion^39^, and rate of lipolysis^40^, support a correlation between adipocyte size and insulin resistance. Consistent with the correlation between adipocyte size and function, *Rreb1* heterozygous mice have smaller adipocytes with increased insulin sensitivity. Human carriers of the *RREB1* variant rs112492319 also have differences in subcutaneous and visceral adipocyte area, with carriers of the T2D-risk allele having increased adipocyte size. This improved insulin sensitivity likely compensates for the detrimental effects of RREB1 loss-of-function on pancreatic beta cell development and function.

Previous studies have shown that RREB1 is required for the generation of brown adipocytes from precursor cells^16,41^. *Rreb1* was significantly upregulated during mesenchymal stem cell differentiation to brown fat in mice^16^. The promoter region of *Rreb1* contains an active super-enhancer bound by the master adipogenic transcription factor PPARγ^16^. Overexpression of *Rreb1* increased thermogenic capacity in brown adipocytes^16^, while knockdown decreased adipogenesis^16,41^. Mechanistically, Rreb1 positively regulates expression of brown fat genes *Ucp1* and *Cidea* by recruiting the H3K27me3 demethylase, Jmjd3, to promoters^41^. Here we show using both human and mouse cellular models that RREB1 loss-of-function decreases expression of pro-adipogenic genes, supporting an additional role for RREB1 in white adipogenesis. Despite a significant decrease in the brown adipocyte gene *Elovl3* in our SGBS knockdown model, we did not measure any significant differences in BAT following *Rreb1* haploinsufficiency *in vivo*.

In conclusion, our work suggests that genetic variation at the *RREB1* locus protects against T2D by impacting progenitor cell differentiation resulting in smaller, more insulin sensitive adipocytes. We have characterized an *in vivo* global heterozygous knockout mouse model, which supports a key role for the adipose tissue in mediating T2D-risk at the *RREB1* locus. Future studies aimed at understanding the tissue-specific contribution of RREB1 will be important to further dissect the genetic contribution of *RREB1* in diabetes.

## Methods

### Animal models and animal care

Global heterozygous C57BL/6N-Rreb1^tm1b(EUCOMM)Wtsi^ knockout mice (homozygotes are embryonic lethal) were obtained from International Mouse Phenotyping Consortium (IMPC) (Sanger, UK)^42–45^. Mice were maintained following UK Home Office legislation and local ethical guidelines issued by the Medical Research Council (Responsibility in the Use of Animals for Medical Research, July 1993). Procedures were approved by the MRC Harwell Animal Welfare and Ethical Review Board (AWERB). Mice were maintained in an animal room with 12h light and dark cycle with a temperature of 21±2°C and 55±10% humidity. Mice were fed *ad libitum* either 1) a standard rat and mouse No. 3 breeding diet (RM3; Special Diets Services, France) containing 11%kcal fat, 27%kcal protein, and 62%kcal carbohydrates; 2) a high-fat diet (HFD; D12492, Research Diets, New Brunswick, NJ, USA) of 60%kcal fat, 20%kcal protein, and 20%kcal carbohydrates; or 3) a low-fat diet (LFD; D12450J, Research Diets, New Brunswick, NJ, USA) of 10%kcal fat, 20%kcal protein and 70%kcal carbohydrates. Phenotyping tests were performed according to EMPReSS (European Phenotyping Resource for Standardised Screens from EUMORPHIA) standardized protocols as described (http://empress.har.mrc.ac.uk/). The cohort of mice used for *in vivo* glucose uptake assays were fed standard chow of 18% calories from fat, 24% calories from protein, and 58% calories from carbohydrates (2018 Teklad Global 18% Protein Rodent Diet). *In vivo* glucose uptake assays were performed per procedures approved by the Institutional Animal Care and Use Committee of the Stanford Animal Care and Use Committee (APLAC) protocol number #32982.

### Body composition

Body composition, including whole body fat mass and lean mass, were measured using Nuclear Magnetic Resonance (EchoMRI-100H). Briefly, mice were restrained in a clear plastic tube before insertion into the instrument for body composition measurements. Body length was measured (in mm) from the tip of the nose to the base of the tail using a ruler. Bone mineral content and bone mineral density were accessed using the II Dual Energy Xray Analyser (DEXA) (PIXImus). An intraperitoneal injection of anesthetic solution (10 µl ketamine/Xylazine per gram of body weight) was given to mice prior to the DEXA procedure. MicroCT analysis was performed using Skyscan 1172 to access tibia tubercular bone. At 38 weeks old, mice were humanely euthanized by cervical dislocation and individual fat depots were dissected and weighed: gonadal WAT (gWAT), mesenteric WAT (mWAT), perirenal WAT (pWAT), inguinal WAT (iWAT)^46,47^.

### Histological analysis

Liver, white adipose tissues (gWAT, gonadal; mWAT, mesenteric; pWAT, perirenal; and iWAT, inguinal subcutaneous), brown adipose tissue were dissected and fixed in Surgipath neutral buffered formaldehyde (Leica). Tissues were embedded with paraffin and were stained with hematoxylin and eosin. Images were captured using the NanoZoomer slide scanner (Hamamatsu Photonics) and adipocyte size was calculated using Adiposoft^48^. Adipocytes were sorted into size ranges. For the HFD and LFD comparisons in both sexes, the size range of each bin differed by 250 µm^2^ increments. For the RM3 initial diet study only males were analyzed and bin sizes increased in 1000 µm^2^ increments. Where relative frequency of adipocyte area data for heterozygotes and wildtype mice converged/diverged, a cutoff size range was determined by visual inspection of the data to allow calculation of area under the curve in the regions of divergence (shown by dotted lines in the Figures). This allowed testing of differences within larger and smaller adipocyte size range. AUC was also calculated for the entire dataset. AUC determination and statistical tests were carried out in GraphPad Prism. Data was evaluated for normality using a combination of D’Agostino and Pearson tests in combination with QQ plots. Depending on normality and equality of variance two tailed t-tests or Mann-Whitney two tailed tests were used.

### Plasma analysis

Plasma insulin was measured using a Mouse Insulin ELISA (Mercodia, 10-1247-01). Adiponectin was measured using a Mouse Adiponectin ELISA (Merck, EZMADP-60K). Leptin was measured using Millipore Leptin ELISA (Millipore, EZML-82k). Plasma liver function markers (ALP, ALT, AST) were measured on an AU400 Automated Clinical Chemistry Analyzer (Olympus).

### Food intake and metabolic rate measurement

For food intake studies, two mice of the same genotype and sex were weaned into a cage at 3 weeks of age. A fine scale was used to measure weekly food weights at the same time as cage changing. Metabolic rate was accessed using Oxymax indirect calorimetry (Columbus Instruments) including oxygen consumption, carbon dioxide production, respiratory exchange ratio (RER), and heat production^49^. Oxygen consumption and carbon dioxide production were normalized to body weight. Heat production (energy expenditure) was determined using the equation heat = CV x VO2, where CV = 3.815 + 1.232 x RER (CV, calorific value based on the observed respiratory exchange ratio)^49^.

### *In vivo* measurements of glucose homeostasis

Intraperitoneal glucose tolerance tests (IPGTT) were performed at 12 and 20 weeks of age. Mice were fasted for 16 hours and were weighed before the test was performed. A blood sample was collected at time zero/baseline from the tail vein using a Lithium-Heparin microvette tube (Sarstedt) after local anesthetic (EMLA cream, Eutectic Mixture of Local Anesthetics, Lidocaine/Prilocaine, AstraZeneca). Mice were intraperitoneal injected with a glucose dose of 2 g/kg body weight (20% glucose in 0.9% NaCl). Blood glucose measurements were performed as above at 30-, 60-, and 120-minutes post glucose injection. Plasma glucose was measured using an GM9 Glucose Analyser (Analox Instruments).

Intraperitoneal insulin sensitivity tests (IPIST) were performed at 16 and 24 weeks of age. Mice were fasted for 5-6 hours and were weighed before the test was performed. A baseline glucose level was measured before mice were injected intraperitoneally with insulin solution. Mice were intraperitoneal injected with 0.5-1.5 IU insulin/kg body weight (0.05-0.15 IU /ml insulin diluted in 0.9% NaCl). Blood glucose measurements were performed at 15-, 30-, 45-, 60-, and 90-minutes post injection.

### Pancreatic islet isolation

The islet isolation procedure was performed as previously described^50^. In brief ice-cold Collagenase XI 0.5 mg/mL solution (Sigma-Aldrich, C7657) was injected into the bile duct and the pancreas was perfused. The pancreas was then removed and incubated at 37°C for 15 minutes with vigorous shaking every 5 minutes. To wash the samples, 0.5% (v/v) fatty acid free BSA solution (Sigma-Aldrich, A8806) was added and the samples were placed on ice for 10 minutes. The washing step was repeated once more before the islets were hand-picked under the microscope.

### *In vitro* glucose stimulated insulin secretion assay (GSIS)

Pancreatic islets were isolated from 13-14 week old female *Rreb1*^+/+^ and *Rreb1*^+/-^ mice (*n* = 4 mice per genotype) and cultured overnight in RPMI-1640 GlutaMax™ (Gibco, 61870036) containing 10% FBS and 1% Pen-Strep. The following day islets were equilibrated in 2 mM glucose in Krebs*-* Ringer HEPES (KRH) buffer containing 0.2% (w/v) fatty acid free BSA (Sigma-Aldrich, A7030) for 1 hour (37°C, 5% CO_2_). Five islets of approximate equal size were hand-picked into each well of a multi-well plate for static GSIS. Islets were treated for 1 hour (37°C, 5% CO_2_) with 5 different secretagogues. At the end of the incubation, supernatant was collected and islets were collected in acid-ethanol solution (75% (v/v) ethanol, 2% (v/v) HCl, 0.1% (v/v) Triton X-100). This process was repeated two more times to achieve 3 technical replicates in assessing insulin secretion per secretagogue condition. Supernatant and islets in acid-ethanol were stored at -20°C before further analysis. Secreted insulin in supernatant samples was measured using the Ultra-Sensitive Mouse Insulin ELISA Kit (Crystal Chem). Islets in acid-ethanol were sonicated for 5 seconds at 40 kHz, centrifuged (16,000 x *g*, 4°C, 15 min) and total protein content measured using the Pierce™ BCA Protein Assay Kit (ThermoFisher Scientific). Three technical replicates were used for each secretagogue condition except for the 20 mM glucose and 500 µM tolbutamide condition, where for one wildtype mouse only two technical replicates were used. Each data point represents the average of the technical replicates.

### Glucose uptake assay

For *in vitro* glucose uptake assays cells were washed with PBS and medium was changed to KRH (50 mM HEPES, 136 mM NaCl, 1.25 mM MgSO_4_, 1.25 mM CaCl_2_, 4.7 mM KCl, pH 7.4) buffer for three hours. Cells were treated with either BSA control or 0.1 mg/mL insulin and left to incubate for 3 hours. After incubation, cells were treated with a mixture of 2-deoxyglucose and 3H-2-deoxyglucose for 10 minutes at 37°C. Cells were then washed twice in cold KRH buffer before being lysed in RIPA lysis buffer for liquid scintillation counting. Data are expressed as raw CPM counts or normalized to total protein. *In vivo* glucose uptake assays were performed as described previously^51^. Briefly, mice were fasted for 2 hours and injected intraperitoneally with ^3^H-2-deoxyglucose at 100 uCi/kg with saline or 1U/kg insulin in a total volume of 120 µL per mouse. After 30 minutes, the mice were euthanized. Wet tissue weights were recorded and then homogenized in 0.1% SDS for liquid scintillation counting. Data are expressed as CPM/mg wet tissue weight.

### SVF isolation and *in vitro* differentiation

Individual mouse adipose depots were carefully excised and placed in PBS. Tissues were cut into 1-2 mm pieces, and transferred into digestion buffer [sterile Hank’s Balanced Salt Solution (Sigma Aldrich, H8264), 0.8 mg/mL (0.08%) collagenase, type II (Worthington Biochemical Corp., LS004174), and 30 mg/mL (3%) bovine serum albumin (Sigma Aldrich, A9418)]. Tissues were incubated for 20-45 minutes with vigorous shaking by hand every 10 minutes. Digestion was considered complete when there were no more visible clumps of tissue. To isolate the SVF fraction, samples were centrifuged for 3 minutes at 300x*g* and the cell pellet was resuspended in growth media (DMEM with 10% FBS and 1% Pen/Strep). The resulting cell suspension was filtered through a 40 µm nylon mesh and plated onto a 10 cm dish. After 24 hours, the attached cells were gently washed twice with PBS before adding fresh growth media. When cells reached 80-90% confluency they were passaged using TrypLE Express (ThermoFisher, 12604013) and were plated at a density of 180,000-200,000 cells/well of 6-well plate for the Oil Red O studies, or 60,000 cells/well for proliferation studies. Twenty-four hours after plating (day 0), media was changed to DMEM with 10% FBS, 125 mM indomethacin, 1 nM T3, 1 µM dexamethasone, 0.5 µM IBMX, and 20 nM human insulin. On day 4, media was changed to DMEM with 10% FBS, 1 nM T3, and 20 nM human insulin. Oil Red O assay was performed at day 7.

### Oil Red O assay

Cells were washed with DPBS twice and fixed with 4% paraformaldehyde for 15 minutes on ice. Cells were then rinsed twice with water and incubated with 60% isopropanol for 5 minutes at room temperature. A 3:2 dilution of Oil Red O filtered solution (0.5 g/100 mL isopropanol; Sigma-Aldrich) was added to cells and incubated for 5 minutes at room temperature. Oil Red O was then rinsed with ddH_2_O three times before addition of a 4% (v/v) IGEPAL® CA-630 (4 mL in 96 mL 100% isopropanol) solution for 10 minutes with gentle shaking on an orbital shaker (Sigma-Aldrich) at room temperature. Lipid content quantification was measured using 100 µL of sample that was transferred into a 96 well plate at an absorbance of 500 nm.

### Click-iT® Plus EdU proliferation assay

Cells were incubated with 10 µM EdU at 37°C for 2 hours. Cells were rinsed with DPBS three times before fixation with 4% PFA for 15 minutes at room temperature. Cells were rinsed twice with 3% (w/v) BSA in PBS and then permeabilized in 0.5% TritonX-100 for 20 minutes at room temperature. Cells were incubated for 30 minutes at room temperature with the Click-iT Plus EdU Cell Proliferation Kit according to manufacturer’s instructions (ThermoFisher Scientific, C10640). Nuclear staining was performed using the NucBlue™ Live ReadyProbes™ Reagent (ThermoFisher Scientific, R37605) and images were captured at 488 nm excitation wavelength by the Zeiss LSM700 confocal microscope..

### SGBS differentiation and gene silencing

SGBS cell line was maintained and differentiated as previously described^27,52^ and experiments were performed using cells between generation 34 and 39. Cells were cultured in DMEM/F12 with 10% FBS, 33 µM biotin (Sigma-Aldrich, B4693), 17 µM pantothenate (Sigma-Aldrich, P5155), and antibiotics (Gibco, 15140-122; 100 IU/mL penicillin and 100 µg/mL streptomycin). For differentiation, 20,000 cells were plated per well of a 12-well plate. Once cells reached near confluence (day 0, approximately three days post plating), cells were washed with PBS before adding DMEM/F12 supplemented with 33 µM biotin, 17 µM pantothenate, antibiotics, 0.01 mg/mL transferrin (Sigma-Aldrich, T2252), 20 nM insulin (Gibco, 12585-014), 100 nM cortisol (Sigma-Aldrich, H0888), 0.2 nM T3 (Sigma-Aldrich, T6397), 25 nM dexamethasone (Sigma-Aldrich, D1756), 250 µM IBMX (Sigma-Aldrich, I5879), and 2 µM rosiglitazone (Cayman, 71740). On day 4, the media was replaced with DMEM/F12 supplemented with 33 µM biotin, 17 µM pantothenate, antibiotics, 0.01 mg/mL transferrin, 20 nM insulin, 100 nM cortisol, and 0.2 nM T3. Media was replaced on day 8 and cells were collected at day 12 for further analysis. Gene silencing was performed according to Lipofectamine RNAiMAX transfection protocol using 50 nM SMART pool (mixture of four siRNAs) on-TARGETplus siRNAs against either non-targeting control (siNT) or *RREB1* (si*RREB1*). Transfection was performed on day -2 or 48-hours before differentiation and knockdown efficiency was assessed at day 0.

### hiPSC differentiation to adipocytes and osteoblasts

*RREB1* wildtype and knockout hiPSC lines were previously generated^21^. The STEMdiff Mesenchymal Progenitor Kit (StemCell Technologies, 05240) was used to generate mesenchymal progenitor cells from *RREB1* wildtype and knockout hiPSC lines. Briefly, hiPSCs were plated on Matrigel-coated plates (Corning, 354230) at a density of 5x10^4^ cells/cm^2^ in mTeSR1 (StemCell Technologies, 85850) supplemented with 10 µM Y-27632 (StemCell Technologies, 72304). Forty-eight hours after plating, media was changed to STEMdiff-ACF Mesenchymal Induction Medium for four days with daily media changes. On day 4, the cells were switched to MesenCult-ACF Plus Medium and passaged accordingly to manufacturer’s protocol on day 6. The cells were then passaged twice weekly at 80% confluency and maintained in MesenCult-ACF Plus Medium. The resulting mesenchymal progenitor cells were cryopreserved using MesenCult-ACF Freezing Medium. MesenCult Adipogenic Differentiation Kit (Human) and MesenCult Osteogenic Differentiation Kit (Human) were used to induce differentiation of hiPSC-derived mesenchymal progenitor cells to adipocytes and osteoblasts, respectively (StemCell Technologies, 05412 and 05465).

### RNA isolation and transcriptomic analyses

For mouse adipose tissues and islets, total RNA was extracted using the Lipid Tissue Mini kit (Qiagen) and the RNeasy Micro kit (Qiagen), respectively, according to the manufacturer’s instructions. RNA concentration was measured using a NanoDrop ND-1000 spectrophotometer (ThermoFisher Scientific). RNA integrity was accessed using the 2100 Bioanalyser (Agilent). Total RNA quantity and integrity were assessed using Quant-IT RiboGreen RNA Assay Kit (Invitrogen) and Agilent Tapestation. Purification of mRNA, generation of double stranded cDNA and library construction were performed using NEBNext Poly(A) mRNA Magnetic Isolation Module (E7490) and NEBNext Ultra II Directional RNA Library Prep Kit for Illumina (E7760L) with our own adapters and barcode tags (dual indexing^53^). The concentrations used to generate the multiplex pool were determined by Picogreen. The final size distribution of the pool was determined using a Tapestation system (Agilent), and quantification was determined by Qubit (ThermoFisher Scientific) before sequencing on an Illumina NovaSeq6000 as 150bp paired end. For SGBS samples, total RNA was extracted using TRIzol according to manufacturer’s instructions (ThermoFisher Scientific, 15596026). Sample QC, mRNA library preparation (poly A enrichment), and RNA-seq (NovaSeq PE150) were performed by Novogene. Reads were mapped to the human genome build GRCh38 or mouse genome build GRCm38 using STAR (v2.7.9a)^54^ with ENSEMBL gene annotations (v101). featureCounts (v2.0.1) was used to determine gene expression levels and differential expression analysis was performed using DESeq2^55^.

### Analysis of enhancer activity and interactome during adipogenesis and osteogenesis

Fastq files from ATAC-Seq during SGBS differentiation were obtained from GSE178796^56^ and processed using the nextflow pipeline (https://nf-co.re/atacseq/2.1.2, version 2.1.2) developed and maintained by the nf-core framework^57^. Briefly, reads were aligned to the hg19 reference genome using BWA^58^ and normalized coverage tracks created using BEDtools^59^. MED1, H3K27ac ChIP-Seq and DNase-seq [GSE113253] were processed as previously described^60^. Data for enhancer capture Hi-C at Day 10 of BM-hMSC-TERT adipogenesis [GSE140782] was processed as previously described^61^. Integration and visualization of genomic data was performed using the python package pyGenomeTracks v3.8^62^.

### Adipocyte area measurements in human donor adipose tissue

Adipocyte area estimates, together with genotype calls based on the Illumina Global Screening bead chip array, were retrieved from the Munich Obesity BioBank (MOBB) to investigate the relationship between fat cell size and *RREB1* genotype in a large cohort of human samples. Briefly, subcutaneous and visceral adipose tissue, together with buffy coat for genomic DNA extraction were collected during elective abdominal surgery. Written informed consent was obtained from all study participants and the study protocol was approved by the ethics committee of the Technical University of Munich (Study No. 5716/13). Hematoxylin and Eosin-stained slides were generated from formalin-fixed paraffin embedded tissue and adipocyte area was determined using a machine learning based approach (Adipocyte U-net) as described earlier^32,63^. A lower bound threshold of 200 µm^2^ and an upper bound threshold of 16,000 µm^2^ was set to remove potential artefacts. Mean adipocyte area per depot and individual was calculated based on a number of 200 unique identified objects and used for all further analyses stratifying between different *RREB1* genotypes.

## Statistical analysis

All data are displayed as means ± s.e.m. Statistical analyses were generated using Prism 10.0.2 (GraphPad). Details of the statistical tests used are indicated in the figure legends. Statistical significance is represented as \**p*<0.05, \*\**p*<0.01, \*\*\**p*<0.001, \*\*\*\**p*<0.0001.

## Data Availability

Sequencing data will be available at the European Genome-phenome Archive (EGA) under study number EGAS50000000253.

## Acknowledgements

We thank the Wellcome Trust Sanger Institute Mouse Genetics Project (Sanger MGP) and its funders for providing the mutant mouse line C57BL/6N-Rreb1^tm1b(EUCOMM)Wtsi^ and the European Mouse Mutant Archive (www.infrafrontier.eu; Repository number EM:14150) partner the Mary Lyon Centre at MRC Harwell from which the mouse line was received. Funding and associated primary phenotypic information may be found at www.sanger.ac.uk/mouseportal and https://www.mousephenotype.org. We thank the UKRI Medical Research Council (MC_U142661184, MC_UP_2201/1) for supporting the mouse studies. NAJK was supported by the Stanford Maternal and Child Health Research Institute (MCHRI) Postdoctoral fellowship. MZ was supported by the K99/R00 NIH Pathway to Independence Award (K99AR081618) from NIAMS. JWK was supported by the NIH through grants: P30 DK116074 (to the Stanford Diabetes Research Center), R01 DK116750, R01 DK120565, R01 DK106236; and by the American Diabetes Association through grant 1-19-JDF-108. KJS was supported by R01DK125260, Stanford Diabetes Research Center P30DK116074, and American Heart Association 23IPA1042031. This work was funded in Oxford and Stanford by the Wellcome Trust (095101, 200837), the UKRI Medical Research Council (MR/L020149/1), and the NIH (UM1-1DK126185). ALG was supported by a Wellcome Trust Senior Fellow in Basic Biomedical Science.

## Author contributions

Conceptualization, R.D.C., A.L.G.; Formal Analysis, H.S., J.H., M.N., H.D.; Investigation, G.Z.Y., N.A.J.K, L.Z., M.Z., K.P.,Y.B., M.R., S.T., V.R., F.A., R.C., A.L.G; Resources, M.W., K.K.M; Data Curation, H.S., J.H., M.N., H.D.; Writing – Original Draft, N.A.J.K, A.L.G.; Writing – Review & Editing, G.Z.Y., N.A.J.K., M.Z., K.P., M.N., S.M., R.D.C., A.L.G.; Visualization, G.Z.Y., N.A.J.K., M.Z., K.P., H.S., M.N., J.H., H.D., and R.D.C.; Supervision, H.H., J.W.K., J.Y.W, S.M., M.C., K.J.S., R.D.C., A.L.G.; Project Administration, R.D.C., A.L.G.; Funding Acquisition, R.D.C., A.L.G.

