## Supplemental Figures for "Loss of *RREB1* reduces adipogenesis and improves insulin sensitivity in mouse and human adipocytes"

### Extended Data Figures

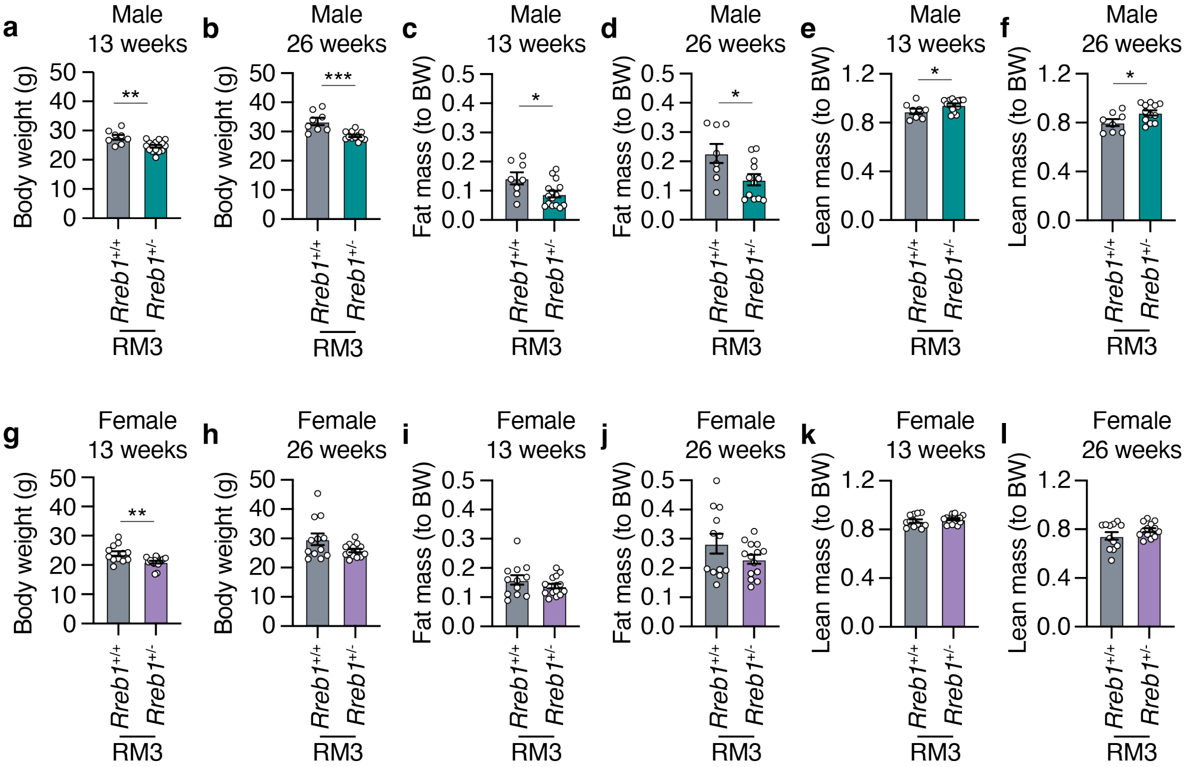


**Extended Data Fig. 1 | Reduced body weight *Rreb1* heterozygous knockout mice on RM3 diet. a,b** Body weight (g) measurements of wildtype (grey) and *Rreb1* heterozygous knockout (green) male mice at (**a**) 13 weeks (*n* = 8-15) and (**b**) 26 weeks (*n* = 8-12) on RM3 diet. **c,d** Fat mass normalized to body weight (BW) of wildtype (grey) and *Rreb1* heterozygous knockout (green) male mice at (**c**) 13 weeks (*n* = 8-15) and (**d**) 26 weeks (*n* = 8-12) on RM3 diet. **e,f** Lean mass normalized to body weight (BW) of wildtype (grey) and *Rreb1* heterozygous knockout (green) male mice at (**c**) 13 weeks (*n* = 8-15) and (**d**) 26 weeks (*n* = 8-12) on RM3 diet. **g,h** Body weight (g) measurements of wildtype (grey) and *Rreb1* heterozygous knockout (purple) female mice at (**g**) 13 weeks (*n* = 12-16) and (**h**) 26 weeks (*n* = 12-14) on RM3 diet. **i,j** Fat mass normalized to body weight (BW) of wildtype (grey) and *Rreb1* heterozygous knockout (purple) female mice at (**i**) 13 weeks (*n* = 12-16) and (**j**) 26 weeks (*n* = 12-14) on RM3 diet. **k,l** Lean mass normalized to body weight (BW) of wildtype (grey) and *Rreb1* heterozygous knockout (purple) female mice at (**k**) 13 weeks (*n* = 12-16) and (**l**) 26 weeks (*n* = 12-14) on RM3 diet. Data are presented as mean ± s.e.m. Statistical analyses were performed by unpaired two-tailed t test. **p*<0.05, ***p*<0.01, and ****p*<0.001.

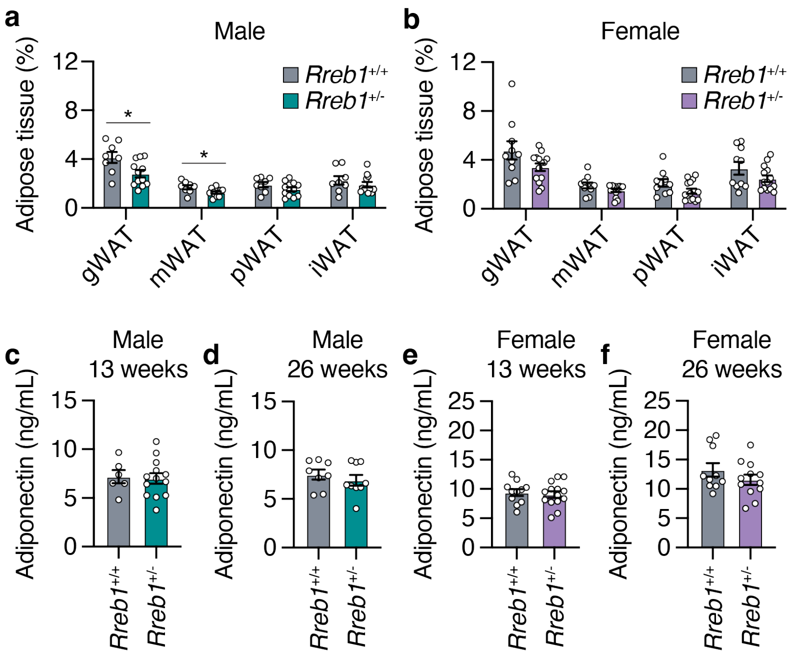


**Extended Data Fig. 2 | *Rreb1* heterozygous knockout male mice have decreased white adipose depot size on RM3 diet. a** Comparisons of gonadal, mesenteric, perirenal, and subcutaneous inguinal white adipose tissue weight (% normalized to body weight) at 38 weeks in control (grey) and *Rreb1* heterozygous knockout (green) male mice on RM3 diet. *n* = 8-12. **b** Comparisons of gonadal, mesenteric, perirenal, and subcutaneous inguinal white adipose tissue weight (% normalized to body weight) at 38 weeks in control (grey) and *Rreb1* heterozygous knockout (purple) female mice on RM3 diet. *n* = 10-14. **c,d** Plasma adiponectin (ng/mL) after an overnight fast at (**c**) 13 weeks and (**d**) 26 weeks for male wildtype (grey) and *Rreb1* heterozygous knockout (green) mice on RM3 diet. *n* = 6-13. **e,f** Plasma adiponectin (ng/mL) after an overnight fast at (**e**) 13 weeks and (**f**) 26 weeks for female wildtype (grey) and *Rreb1* heterozygous knockout (purple) mice on RM3 diet. *n* = 10-14. Data are presented as mean ± s.e.m. Statistical analyses were performed by unpaired t test. *$p<0.05$.

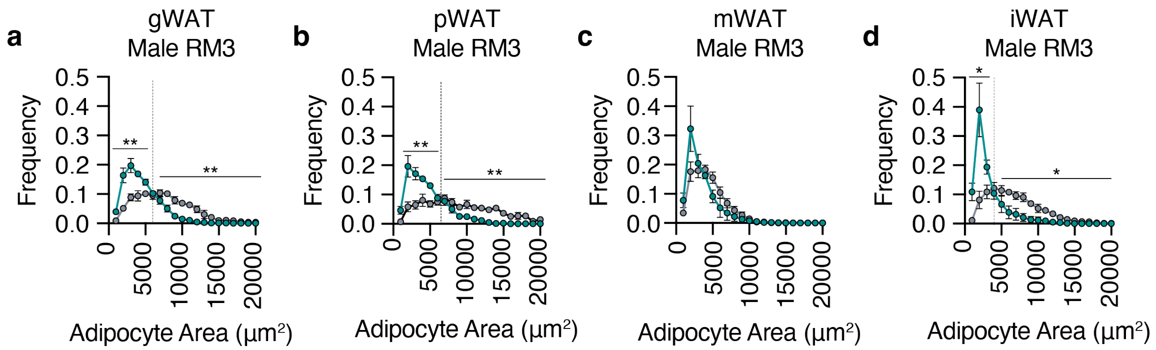


**Extended Data Fig. 3 | Tissue-specific differences in adipocyte size of white adipose tissue in *Rreb1*$^{+/-}$ mice on RM3 diet. a-d** Quantification of adipocyte area (μm$^2$) within (**a**) gonadal visceral (gWAT), (**b**) perirenal (pWAT), (**c**) mesenteric (mWAT), and (**d**) inguinal subcutaneous (iWAT) fat depots from male *Rreb1*$^{+/+}$ and *Rreb1*$^{+/-}$ mice fed a RM3 diet at 38 weeks of age. Dotted line denotes the interval where the dataset converges. **a**, *n* = 3, 4; **b**, *n* = 3, 4; **c**, *n* = 3, 3; and **d**, *n* = 3, 5 *Rreb1*$^{+/+}$ and *Rreb1*$^{+/-}$, respectively. Data are presented as mean ± s.e.m. Statistical analyses were performed by unpaired t test of the area under the curve for adipocytes (**a**) <6000 and >6000 μm$^2$ or (**b**) <6000 and >7000 μm$^2$, or (**c**) <4000 and >4000 μm$^2$ or (**d**) <4000 and >4000 μm$^2$. $^*$p<0.05, $^{**}p$<0.01 and $^{****}p$<0.0001.

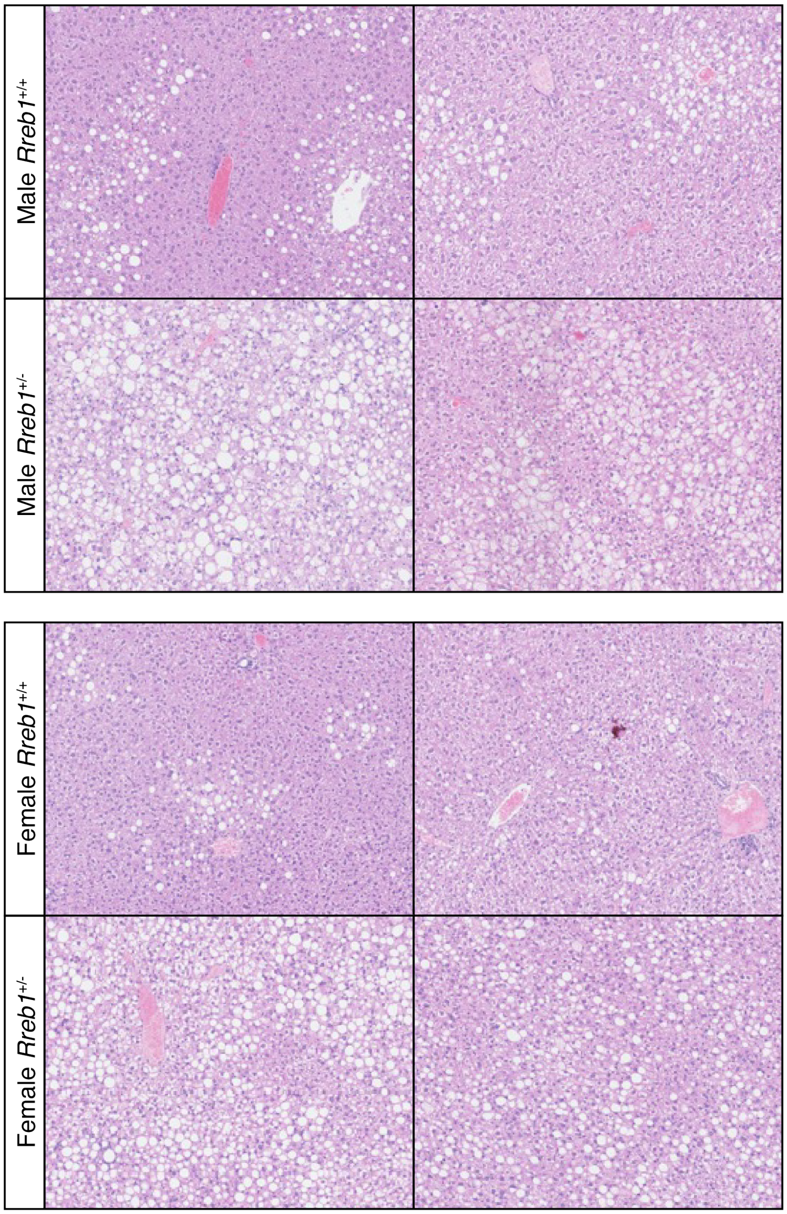


**Extended Data Fig. 4 | Increased lipid droplet formation in liver of *Rreb1*$^{+/-}$ mice.** Representative images of lipid droplet formation in livers of male and female *Rreb1*$^{+/+}$ and *Rreb1*$^{+/-}$ mice on HFD diet.

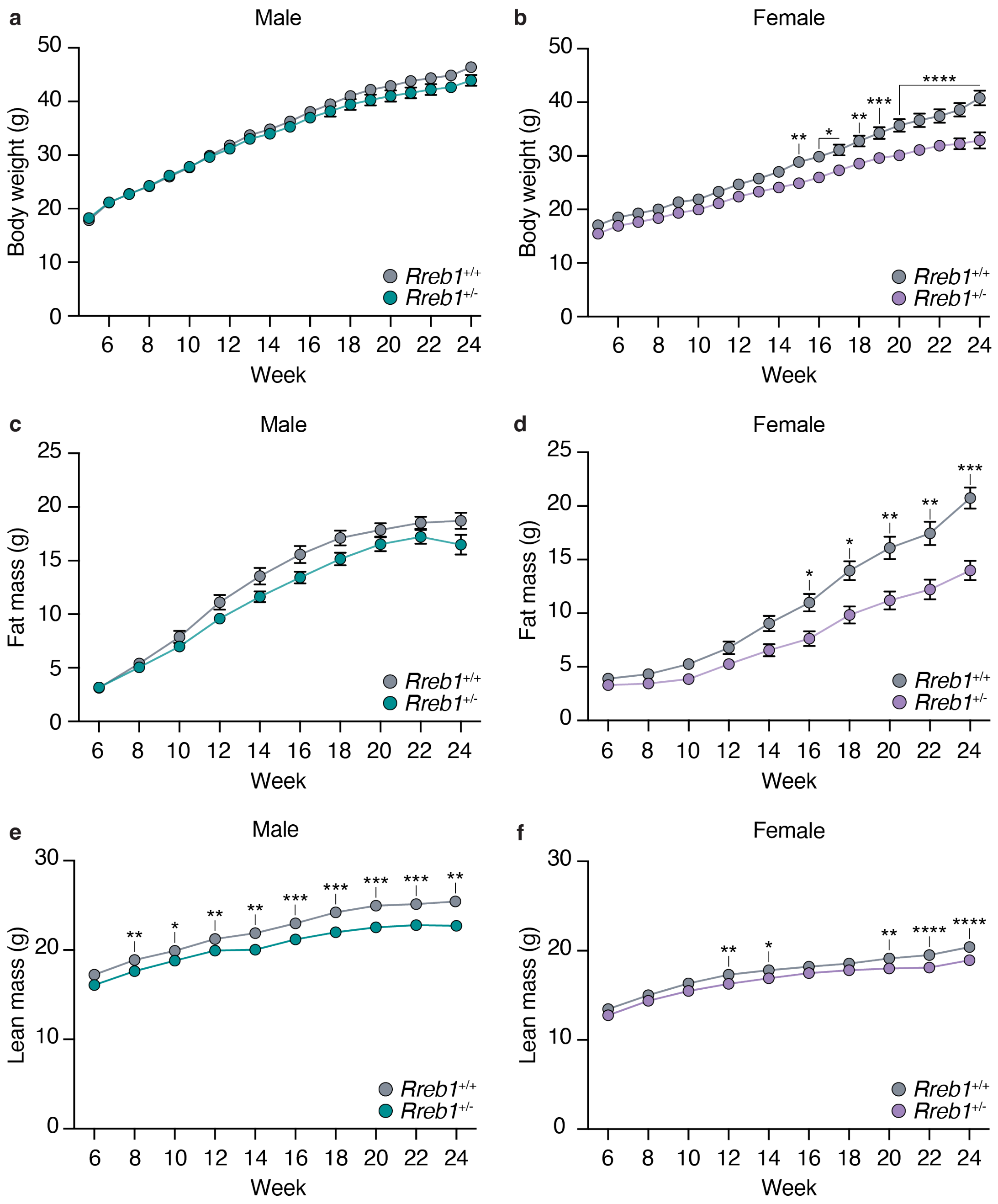


**Extended Data Fig. 5 | Body composition measurements of mice used for food intake studies. a,b** Weekly measurements of body weight (g) of (**a**) male and (**b**) female on HFD. **c,d** Comparisons of fat mass (**c**) male and (**d**) female mice on HFD. **e,f** Comparisons of lean mass (**e**) male and (**f**) female mice on HFD. **a**, $n$ = 19, 24; **b**, $n$ = 20, 20 except for 24 weeks when $n$ = 18,10; **c**, $n$ = 21, 22; **d**, $n$ = 20, 20; **e**, $n$ = 21, 21-22; and **f**, $n$ = 20, 20 *Rreb1*$^{+/+}$ and *Rreb1*$^{+/-}$, respectively. Statistical analyses were performed by two-way ANOVA or in the case of (**b, e**) a mixed effects analysis due to missing data at 24 weeks in (**b**) or two heterozygous very low outliers removed in (**e**), and Bonferroni's (**a,c,d,f**) or Sidak's (**b,e**) multiple comparisons test.

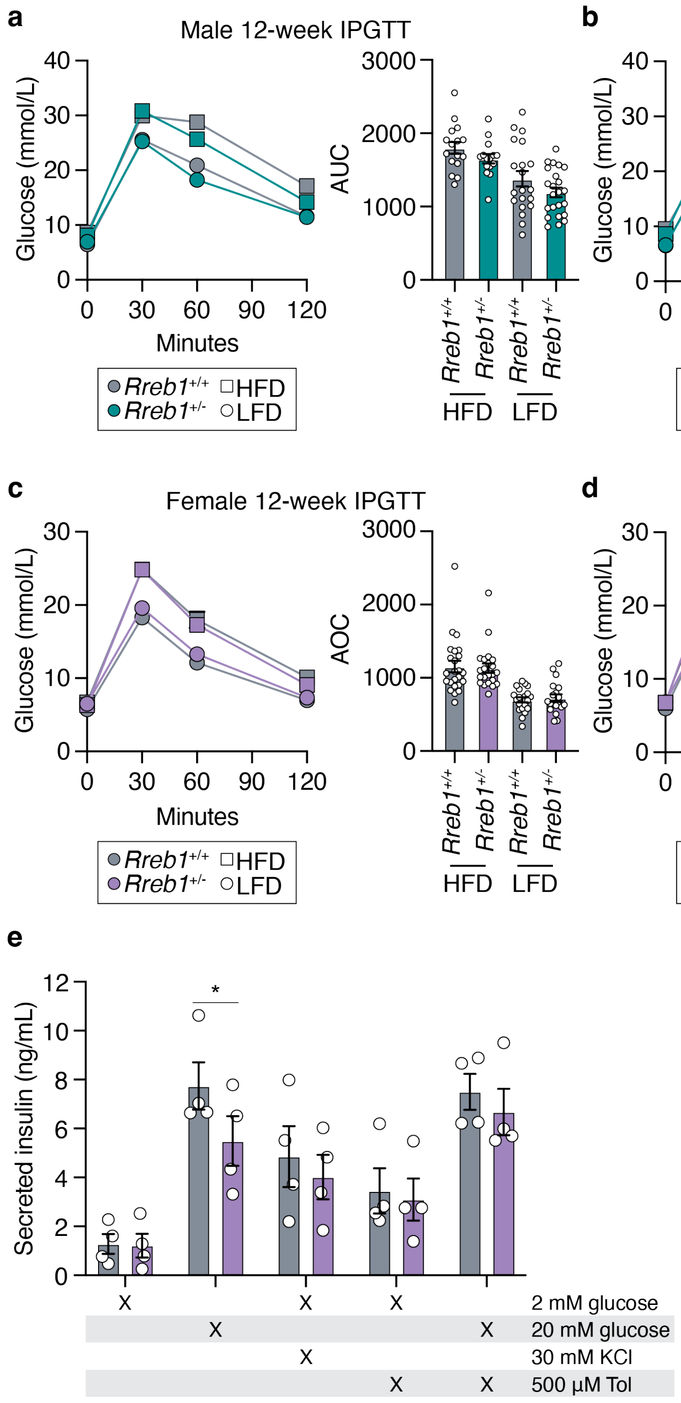


**Extended Data Fig. 6 | Effect of RREB1 on glucose homeostasis and glucose-stimulated insulin secretion. a,b** IPGTT on wildtype and *Rreb1*$^{+/-}$ male mice on HFD and LFD at (**a**) 12 weeks and (**b**) 22 weeks of age. **c,d** IPGTT on wildtype and *Rreb1*$^{+/-}$ female mice on HFD and LFD at (**c**) 12 weeks and (**d**) 22 weeks of age. **e**, Islets from female wildtype and *Rreb1*$^{+/-}$ mice were preincubated in 2 mM glucose for one hour and then treated with either 2 mM glucose, 20 mM glucose, 2 mM glucose and 30 mM potassium chloride (KCl), 2 mM glucose and 500 µM tolbutamide (Tol), or 20 mM glucose and 500 µM Tol for one hour. **a**, HFD: *n* = 16, 15, LFD: *n* =

20, 23; **b**, HFD: *n* = 16, 15, LFD: *n* = 20, 22; **c**, HFD: *n* = 24, 22, LFD: *n* = 21,19; **d**, HFD: *n* = 24, 22, LFD: *n* = 20,19 *Rreb1*$^{+/+}$ and *Rreb1*$^{+/-}$, respectively; and **e**, *n* = 4. Data are shown as mean ± s.e.m. Statistical analyses were performed between genotypes using a Brown-Forsythe and Welch ANOVA tests (**a**) or one-way ANOVA (**b, c**, **d**) or (**e**) using two-way ANOVA using Bonferroni multi-comparison test. In the case of (**c**) the data was Y=log(Y) transformed before analysis. *$p$<0.05; **$p$<0.01.

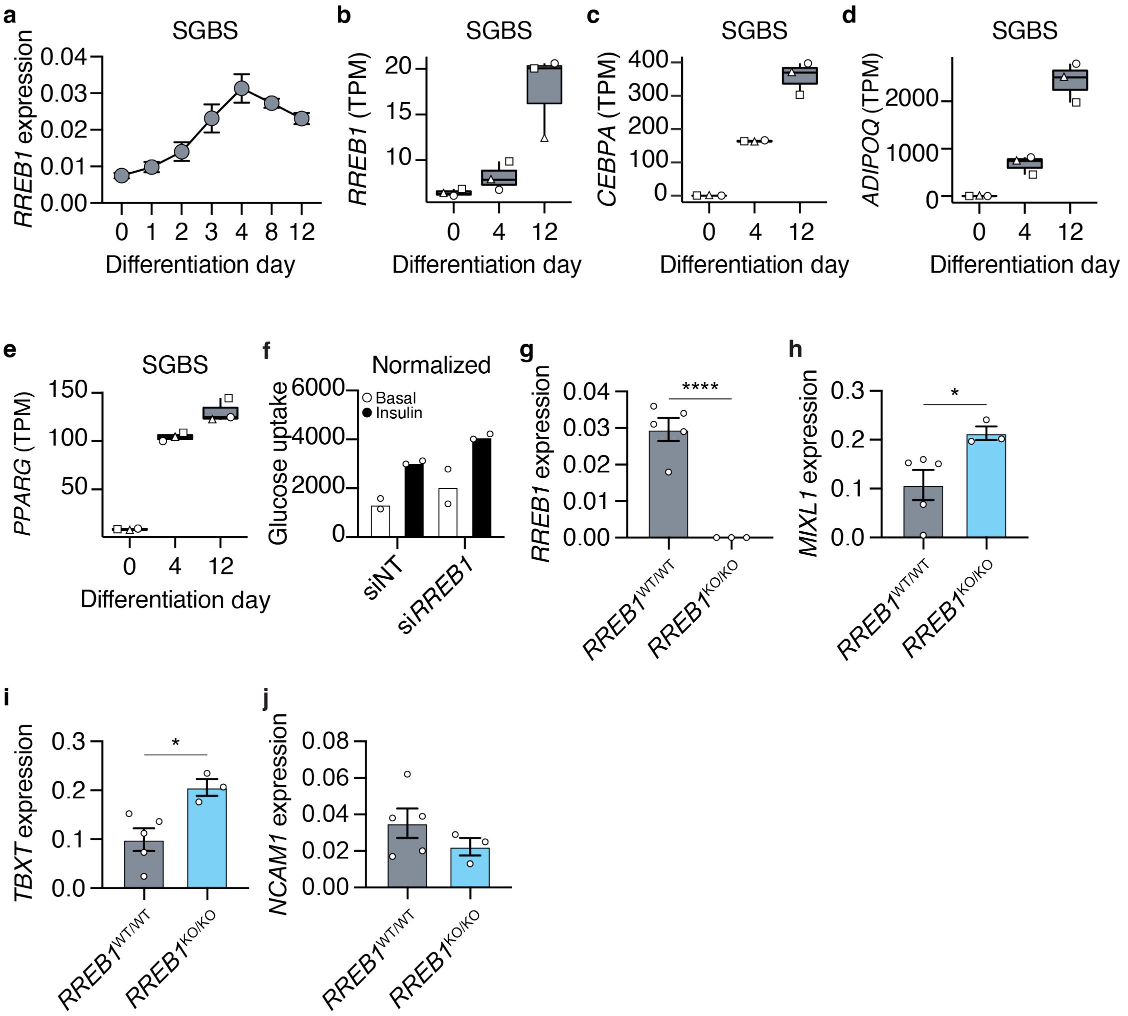


**Extended Data Fig. 7 | RREB1 loss-of-function impacts human *in vitro* adipocyte differentiation. a** *RREB1* transcript expression during SGBS differentiation to adipocytes normalized to *PPIA*. **b,c,d,e** Transcript expression in transcripts per million (TPM) of (**b**) *RREB1*, (**c**) *CEBPA*, (**d**) *ADIPOQ*, and (**e**) *PPARG* during SGBS differentiation to adipocytes. $n = 3$; each replicate is represented by a different symbol. **f** Glucose uptake of SGBS cells at day 12 of *in* vitro differentiation, normalized to total protein content. $n = 2$. SGBS were treated with siNT and si*RREB1* 48 hours before differentiation. **g** *RREB1* transcript expression normalized to *PPIA* in human induced pluripotent stem cell (hiPSC)-derived mesodermal cells. $n$ = 3-4. **h,i,j** hiPSC wildtype (*RREB1*$^{WT/WT}$) and *RREB1* knockout (*RREB1*$^{KO/KO}$) cells were differentiated to mesodermal cells. The expression of mesoderm markers genes (**h**) *MIXL1*, (**i**) *TBXT*, and (**j**) *NCAM1* were measured by qPCR and normalized to *PPIA*. $n$ = 3-5. Data are shown as mean ± s.e.m. Statistical analyses were performed using an unpaired t-test. *$p<0.05$; ****$p<0.0001$.

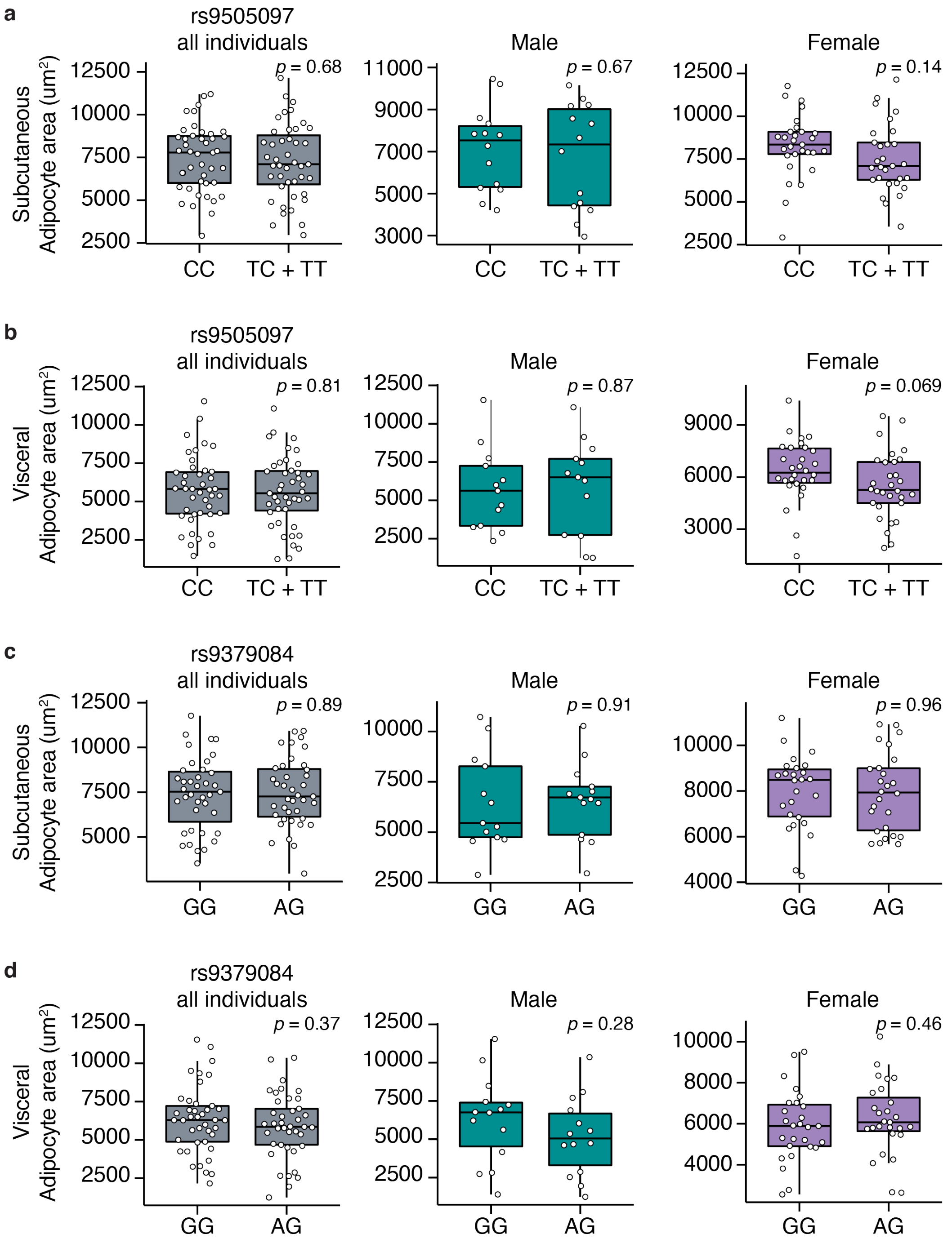


**Extended Data Fig. 8 | No difference in human adipocyte area in *RREB1* rs9505097 and rs9370984 variant carriers. a,b** Adipocyte area of (**a**) subcutaneous and (**b**) visceral fat from human donors carrying rs9505097 variants. **c,d** Adipocyte area of (**c**) subcutaneous and (**d**) visceral fat from human donors carrying rs9370984 variants. Left graph: all individuals (pooled and matched); middle graph: males; right graph: females. Data are presented as mean ± s.e.m. Statistical analyses were performed by unpaired t-test. Individual *p* values are labelled in each graph.

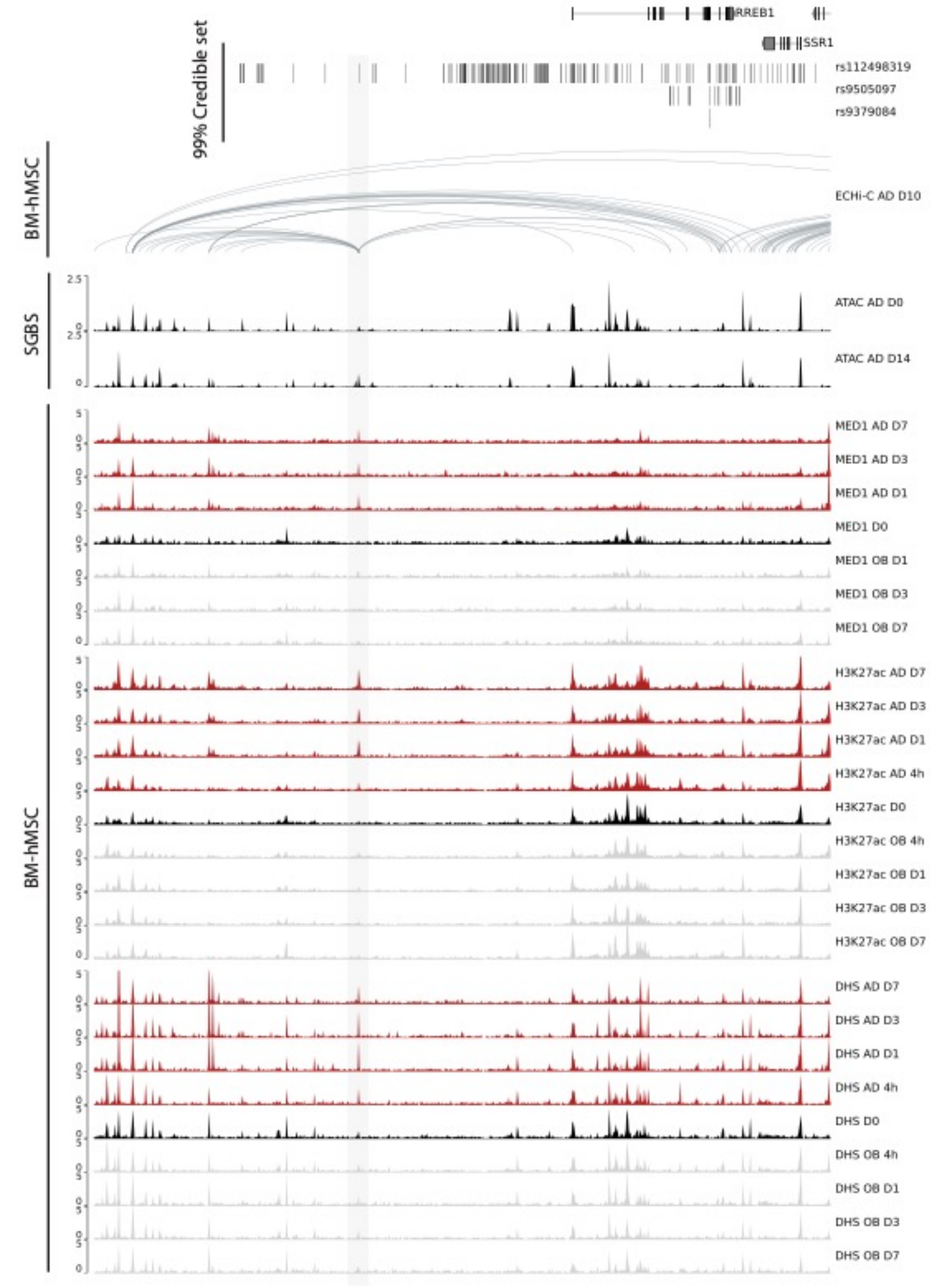


**Extended Data Fig. 9 | Visualization of overlap between regulatory elements and credible-set coverage of variants in the *RREB1* locus during adipogenesis and osteogenesis.** From top: *RREB1* locus (chr6:6667710-7255917) and the 99% credible set of selected variants; significant interactions (FDR < 0.01) from Enhancer Capture Hi-C at day 10 of BM-hMSC-TERT4 differentiated to adipocytes; normalized coverage tracks for ATAC-Seq in differentiating SGBS cells; and normalized coverage tracks of H3K27ac and MED1 Chip-seq, and DNase-Seq in differentiating BM-hMSC-TERT4 cells.

**Extended Data Tables**

**Extended Data Table 1:** Differentially expressed genes in adipose tissue of *Rreb1*$^{+/-}$ mice.

**Extended Data Table 2:** Enriched GO terms from differentially expressed genes in adipose tissue of *Rreb1*$^{+/-}$ mice.

**Extended Data Table 3:** Differentially expressed genes between day 0 and day 12 of SGBS cells differentiated to adipocytes.

**Extended Data Table 4:** Differentially expressed genes in si*RREB1* treated SGBS cells at Day 5 of differentiation to adipocytes.

**Extended Data Table 5:** Enriched GO terms from differentially expressed genes in si*RREB1* treated SGBS cells at Day 5 of differentiation to adipocytes.
